# Conformational stability and order of Hoogsteen base pair induced by protein binding

**DOI:** 10.1101/2023.05.08.539938

**Authors:** Kanika Kole, Aayatti Mallick Gupta, Jaydeb Chakrabarti

## Abstract

Several experimental studies have shown that in the presence of proteins, the Hoogsteen (HG) base pair (bp) becomes stabilized. The molecular mechanism underlying this stabilization is not well known. This leads us to use all-atom molecular dynamics simulation to examine the stability of the HG bp in duplex DNA in both the absence and presence of proteins. We use conformational thermodynamics to investigate the stability of a HG bp in duplex DNA at the molecular level. We also compute the changes in the conformational free energy and entropy of DNA when DNA adopts a HG bp in its bp sequence rather than a Watson–Crick (WC) bp in both naked DNA and protein-bound DNA complex. We observe that the HG bp and the entire DNA duplex conformation are stabilized and ordered in the presence of proteins. Sugar-phosphate, sugar-base, and sugar-pucker torsion angles play key roles in stabilizing and ordering the HG bp in the protein-bound DNA complex.

## 1. Introduction

In double helical DNAs, bp often adopts other geometries than the conventional WC base pairing. HG base pairing is an alternative base-pairing scheme for DNA double helices^1-2^ where the purine is flipped ‘upside-down’ such that the 5-ring of the purine forms a hydrogen bond (H-bond) to the pyrimidine, rather than the 6-ring. The switch from the canonical WC to the non-canonical HG orientation substantially modifies the chemical environment around the bp, which has major implications for DNA−protein recognition^3-4^, damage repair^5-6^, and replication^7-8^. It is in general difficult to analyze HG bp in protein-bound DNA structures by X-ray crystallography due to unclear electron density and by nuclear magnetic resonance (NMR) spectroscopy due to inadequate chemical shift dispersion. As a result, the stability of HG bp in the presence of particular proteins or ligands is far from understood.

HG bp occurs in regions of DNA that are highly distorted by bound protein^9^. The X-ray structure of TATA-binding protein (TBP) bound to DNA reveals a protonated G-C^+^ HG bp in the region of DNA under winding and intercalation by TBP’s phenylalanine side chain^10^. Furthermore, the X-ray crystal structure of DNA in contact with the p53 tumour suppressor proteins exhibits two consecutive A-T HG bps. Wolberger and co-workers report a single A-T HG bp within an undistorted B-form DNA duplex (PDB id-1K61) in the X-ray structure of MATα2 homeodomain bound to DNA^11^. The crystal structure contains 21 fragments of duplex DNA attached to four MATα2 homeodomains. The DNA shows two binding sites for the four MATα2 homeodomains, two homeodomains, α2A and α2B bind at the specific binding sites, while the other two homeodomains, α2C and α2D bind to DNA non-specifically. Earlier MD simulation studies show that the formation of HG bp in the 1K61 system is more strongly influenced by the non-specifically bound protein α2D^11^. However, there has yet not been any attempt to understand the microscopic mechanism underlying the protein induced stability of the HG bp. Here we study this system to focus on the microscopic mechanisms to stabilize the HG bp in the 1K61 system in the presence of proteins. This system is particularly simple since there is no additional distortion in the DNA structure.

The changes in conformational stability and order in bio-molecular systems are quantified in terms of conformational free energy and entropy changes derived from the fluctuations of microscopic conformational variables^12-14^. Negative changes in conformational free energy and entropy indicate structural stability and ordering, respectively, in a given structure with respect to a reference structure, while positive changes indicate conformational destabilization and disorder respectively. Although originally developed for proteins, the conformational thermodynamics calculations are extended to DNA-protein systems as well^12^. The fluctuations of microscopic conformational variables can be captured via long molecular dynamics (MD) simulation from the changes in fluctuations of microscopic conformational variables of the DNA duplex. The microscopic conformational variables of DNA, described in SI, are: (i) inter-bp step parameters: shift (D_x_), slide (D_y_), rise (D_z_), tilt (τ), roll (ρ) and twist (ω) (ii) intra-bp parameters: stagger (S_x_), shear (S_y_), stretch (S_z_), buckle (κ), propeller (π) and opening (σ) (iii) sugar-phosphate and sugar-base backbone torsion angles: α, β, γ, δ, ε, ξ and χ, and (iv) sugar-puckers: ν0, ν1, ν2, ν3 and ν4.

We study three different systems: (1) WC, a model system of DNA, (2) HG, HG bp containing DNA, and (3) HGP, protein-bound HG bp containing DNA. We compute conformational thermodynamics data of the DNA: (i) in the HG system and (ii) in the protein-bound HGP system, both with respect to the WC system. We calculate the total changes in conformational free energy and entropy for entire DNA systems. We observe that the flexibility of the bare DNA duplex is increased by the HG bp. In the presence of a specific protein α2B and a non-specific protein α2D, the whole DNA duplex becomes stabilized and ordered. The backbone torsion angles of bps fluctuate less to stabilize and organize the HG bp as well as the entire DNA duplex in the protein-DNA complex.

## 2. Methods

### 2.1 System preparations

The HG and HGP systems used for all-atom MD simulations are 15 bp DNA fragments or DNA with one specifically bound, α2B and one nonspecifically bound, α2D homeodomain taken from PDB ID 1K61. The numbering of phosphate, sugar, and bases as taken in our study is shown in Fig. 1(A). The A_7_-T_26_ bp in the molecular structure is a HG bp. In the HGP system, this HG bp is found in the contact overlapping sites of a specifically (α2B) and a non-specifically (α2D) bound homeodomain. So, the DNA base sequence for each structure is 5’C_2_ATGT<u>A</u>ATTCATTTA_16_3’ in the 5’-3’ direction, where the underlined A_7_ forms HG bp with the complementary base T_26_ in the 3’-5’ direction. The rest of the bases form WC pairing with the complementary sequence.

**Fig. 1:**
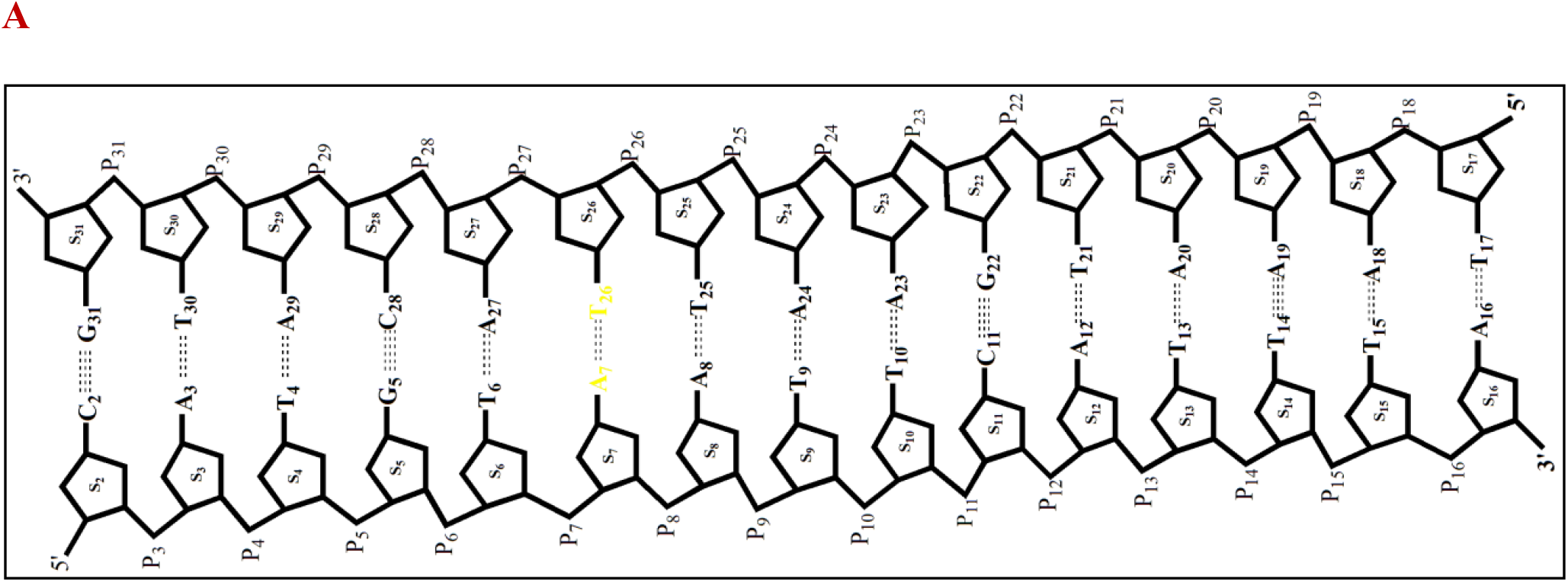
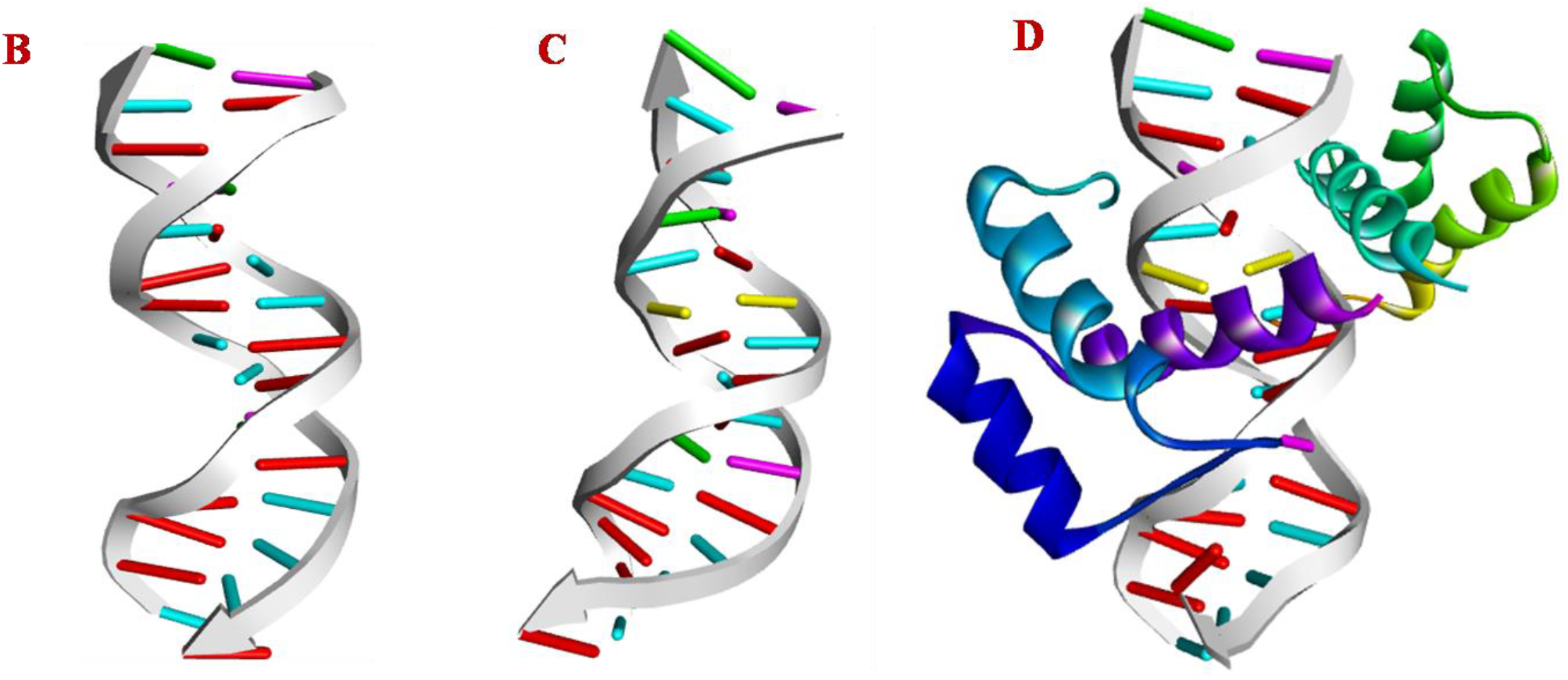
(A) Notations and numbering of DNA bps. Snapshot at 500 ns time span of (B) WC (C) HG (D) HGP systems. HG bp is shown in yellow.

The model structure (WC) of the DNA double helix for the same sequence containing all WC bps has been generated from the fiber-diffraction model of B-DNA^15^ using the SCFBio server (http://www.scfbio-iitd.res.in).

### 2.2 MD simulations

We use the force-field parameters of the Amberff14SB_OL15^16-17^ for DNA and proteins. The GROMACS 2018.6^18^ package standard protocol is used to prepare the systems. The system for simulation is solvated in a cubic water box with a box size of 8.1×8.1 ×8.1nm^3^, containing TIP3P water molecules. The periodic boundary conditions are imposed in all directions. To minimize the end effect, the 3′ end of the DNA strand is connected to its periodic image along the z axis (image patch), to mimic an infinite DNA chain^19^. The system is neutralized by adding the required number of sodium (Na^+^) and chloride (Cl^−^) ions. Subsequently, the system is energy minimized using the steepest descent method^20^. Then all-atom molecular dynamics (MD) simulation is carried out at 300 K and 1 atmospheric pressure in an isothermal-isobaric (NPT) ensemble starting from the energy minimized structure. Berendsen thermostat^21^ is used to maintain a constant temperature, and the pressure is controlled by Parrinello-Rahman barostat^22^. The Lennard-Jones and the short-range electrostatic interactions are truncated at 10 Å. We employ the particle mesh Ewald summation method for long-range electrostatic interaction with 1 Å grid spacing and 10^−6^ convergence criterion. The LINCS constraints are applied to all bonds involving hydrogen atoms. The simulation time step of 2 fs is used for all calculations. We perform MD simulations for the systems (a) WC, (b) HG and (c) HGP, for 1μs. To make the ensembles comparable, we adjust the total number of water molecules so that the numbers of total atoms are the same (N=51068) in all the cases. Various analyses are carried out on the equilibrated portion of the trajectories with tools in GROMACS to examine the system properties. The NUPARM with BPFIND software is used to compute various structural parameters of DNA at intervals of 10 ps^23^.

### 2.3 Structural parameters of the DNA bps

We compute the DNA structural parameters for each of the bp for each system. We divide the equilibrated part of the trajectory into *N*_*w*_ (=10) windows with an equal number of conformations from equilibrated conformations and calculate the averaged value of each parameter, *θ* over each of the windows. We denote the mean value of a parameter, *θ* in the S-th system (S=WC, HG and HGP) by < *θ*^*S*^ >. The error (±0.5Δ*θ*^*S*^) in the mean is given by 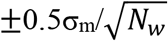, where σ_m_ is the standard deviation of the mean values over the windows. The pseudorotational phase angle P, which is calculated, based on endocyclic torsion angles (ν0, ν1, ν2, ν3 and ν4) inside the ribose sugar ring, is described in SI^24^. All the structural parameters of bps are calculated, ignoring the terminal bps at each end of the DNA duplex.

### 2.4 Conformational thermodynamics

We compute the histograms of all the microscopic variables using the equilibrated trajectory of WC, HG and HGP systems. The normalized probability distribution of any conformational variable, θ in WC, HG, and HGP systems, are given by H^HG^(θ), H^WC^(θ) and H^HGP^(θ), respectively. A detailed description of the histogram-based method (HBM) for calculating the conformational thermodynamics is reported^25-26^.

The changes in free energy of any degree of freedom, θ for each bp of the HG system with regard to its corresponding bp in the WC system and each bp of the HGP system with respect to each bp of WC system are defined as,

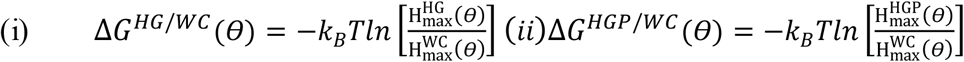

Where “max” represents the maximum of the histogram and *k*_*B*_ is the Boltzmann constant. The change in conformational entropy of a given conformational variable, θ is evaluated from Gibbs formula,

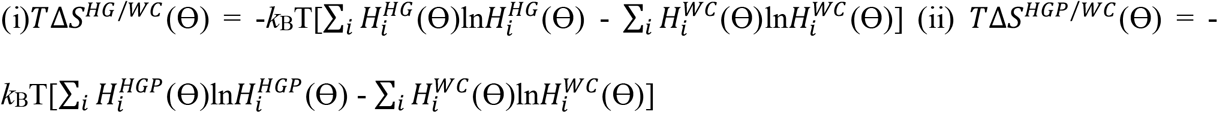

Here 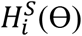 stands for the i^th^ bin of the histogram for θ in the S^th^ case (=WC, HG, HGP).

### 2.5 Interfacial interactions in the HGP system

The interface is identified when two atoms, one from DNA bases and the other from protein (α2D or α2B) residues, are within 0.6 nm of each other. The calculation is performed over the equilibrated portion of the HGP simulated ensemble’s trajectory.

Distance and angle criteria are used to characterize the H-bonds. For H-bonds, the distance between the donor (D) and acceptor (A) atoms is ≤ 0.35 nm and donor-hydrogen-acceptor 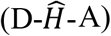 antecedent angle cut off is ≥150^0^. The distance between any of the nitrogen atoms of basic residues of protein and any of the oxygen atoms of the phosphate group of the 5’-3’ strand is used to compute the salt bridge and the cutoff angle conditions are the same as for hydrogen bond interactions. Similarly, the distance and angle between any nitrogen atom from a protein basic residue and any oxygen atom from the phosphate group of the 3’-5’ strand are used to explore the salt bridge interaction for the 3’-5’ strand. If the distance is ≤ 0.4 nm and the angle is ≥ 150^0^, the pair is considered a salt bridge. Ionic species belonging to the DNA base and the protein residue at the interface are considered for electrostatic interactions (attractive charge).When the distance between oppositely charged atoms is less than 0.56 nm, the pair is counted for electrostatic interaction. We utilize GROMACS H-bond analysis module^27^ for H-bonds and Discovery studio software^28^ for salt bridge and electrostatic interaction to define the cutoff criterion.

## 3. Results and Discussions

The WC and HG systems contain 15 bps of DNA alone, whereas the HGP system contains DNA with a specifically bound α2B and a nonspecifically bound α2D homeodomain proteins. A_7_-T_26_ forms a HG base pairing in both HG and HGP systems. The equilibration of each system is ensured by the saturation of the root mean square deviation (RMSD) with time (SI Fig. S2(A)). We consider the equilibrated part of the trajectory, 200 ns-1μs for further analysis. The equilibrated snapshots at 500 ns time span of the WC, HG, and HGP systems are shown in Figs. 1(B), (C), and (D), respectively. The initial snapshot and equilibrated snapshot at 1μs time span of each system are shown in SI Fig. S2.

### 3.1 Conformations in different systems

First, we check the DNA conformations in various cases. SI Tables S1-S3 and S4-S6 show the averages with the errors for six bp step parameters and six intra-bp parameters for each bp in each system. These results show that the average value of each parameter corresponds well to the B-DNA data^29^. The mean value and the error of the mean value of the sugar phase angle, P, for all systems are tabulated in SI Table S7. We observe that the sugar conformation of each base in each system is predominantly C2’-endo like (137^0^-194^0^)^30^, though some transition to C3’-exo (194^0^-216^0^) pucker is also found at the sugars linking to A_7_ and T_26_ bases of the HG and the T_9_ base of the HGP system but the frequencies of these are quite small.

Available data suggest that the transition from the WC to HG geometry requires about a 180^°^ rotation of the base along the bond connecting the base to the sugar, known as the glycosidic bond. The WC N6-H--O4 H-bond is retained in A-T HG bp whereas the other WC N1--H-N3 H-bond is replaced by N7--H-N3 H-bond. In addition, the formation of HG-type H-bonds requires that the two bases come into closer proximity, thus shortening the C1’ –C1’ distance by about 2 Å relative to WC bp. The HG pairing for A-T is sketched in SI Fig. S1. We check the HG base pairing criteria and compare them with literature values^31-35^. We consider the specific hydrogen bonding in A_7_-T_26_. SI Table S8 displays the mean values along with the error of the mean of these parameters (*d*_*D*−*A*_ and 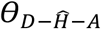) for all systems. We observe that H-bonds form between N1A_7_ (donor) and N3T_26_ (acceptor) in the WC system for A_7_-T_26_ bp which is consistent with the known hydrogen bonding pattern. On the other hand, H-bonds are observed between N7A_7_ (donor) and N3T_26_ (acceptor) in both HG and HGP systems.

The mean and the error associated with the mean of C1’ –C1’ distance (*d*_*C*1′_) for T_6_-A_27_, A_7_-T_26_ and A_8_-T_25_ bps for all the systems are shown in SI Table S9. The HGP system shows the shortest *d*_*C*1′_ in A_7_-T_26_. Apart from A_7_-T_26_ bp, the *d*_*C*1′_ at nearby T_6_-A_27_ and A_8_-T_25_ bps of HG and HGP systems differ from the WC system. No significant changes in *d*_*C*1′_ are observed for the remaining bps. SI Table S9 shows the mean of the neighboring P-P distance (*d*_*p*_) and the error associated with the mean. *d*_*p*_ close to A_7_-T_26_ bp is shortened in both HG and HGP systems (SI Table S9).

The glycosidic torsion angle χ, which characterizes the relative base/sugar orientation in DNA duplexes, is defined by atoms O4′-C1′-N9-C4 and O4′-C1′-N1-C2 in purines and pyrimidines, respectively. We calculate the mean and the error of the mean of χ for each base. The HG and HGP systems show syn *χ* torsion values in the A_7_ base which are consistent with the HG base pairing. The value of χ at complementary T_26_ and adjoining A_8_ bases shows minor variations in HG and HGP systems compared to the WC system (SI Table S10). In addition to T_26_ and A_8_ bases, the χ value for the adjoining T_6_ base in the HGP complex shows a little deviation from the WC conformer. The χ torsion angle does not differ much between systems in the remaining bases. We also observe that α and γ torsions are in the usual gauche^-^/gauche^+ 36^ conformation about base A_7_ for WC and HG systems (SI Table S11). However, in HGP, they follow the unusual gauche^+^/gauche^-^ conformation. α and γ torsion angles of the complementary T_26_ base remain unchanged for all systems. This result is in agreement with the experimental observations^11^.

In the HGP system, for each DNA base, we identify the residues of the protein that form strong contact with DNA bases, as detailed in SI Table S12. As, A_7_-T_26_ bp creates HG bp, we examine in detail the interface interaction for A_7_-T_26_ bp. SI Fig.S3 is a snapshot of active protein residues at the interface of HG bp. We note that protein α2D residues ARG132, GLY133 and GLN175 create interface with base A_7_ of DNA (SI Table S13). These three residues form H-bonds with A_7_ at the interface. Moreover, residue ARG132 is involved in forming salt bridge as well as electrostatic contact between the N atom of ARG132 and the O1P atom of A_7_. On the other hand, the specific protein α2B residues HIS134, PHE136, and TRP179 create H-bonds with T_26_ (SI Table S14). In the 3’-5’ strand, there is no salt bridge or electrostatic contact between T_26_ and the residues of α2B.

### 3.2 Fluctuations in conformational variables

The conformational fluctuations are quantified in terms of the distribution, *H*^*S*^(*θ*_*R*_), (S=WC, HG and HGP) of a microscopic conformational variable, Θ pertaining to base, *R* over the equilibrated trajectory. Each peak of the histogram represents the most probable value for the given variable. The larger width of the histogram peaks denotes more conformational flexibility. We illustrate below a few cases (χ, ε, ν0 and ν3) of microscopic conformational degrees of freedom that exhibit major changes in the A_7_-T_26_ (HG) bp forming region of each system. Fig. 2 shows the distributions of sugar-phosphate (ε), sugar-base (χ) and sugar-pucker (ν0, ν3) torsion angles for A_7_ and T_26_ bases. 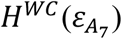 and 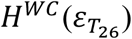 are multi-peaked, as shown in Figs. 2(A) and 2(B). On the other hand, 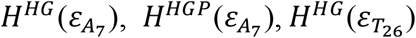 and 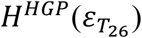 are all double peaked, showing a decrease in flexibility of the HG and HGP systems compared to the WC system for ε. Fig. 2(C) shows that 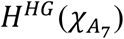 is distributed over a different range compared to 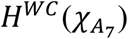. The height of the peak in 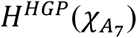 is higher than that of 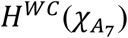. Again, the distributions of 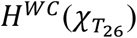 and 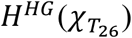 are almost identical (Fig. 2(D)) but 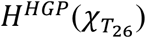 has a sharper peak.

**Fig 2:**
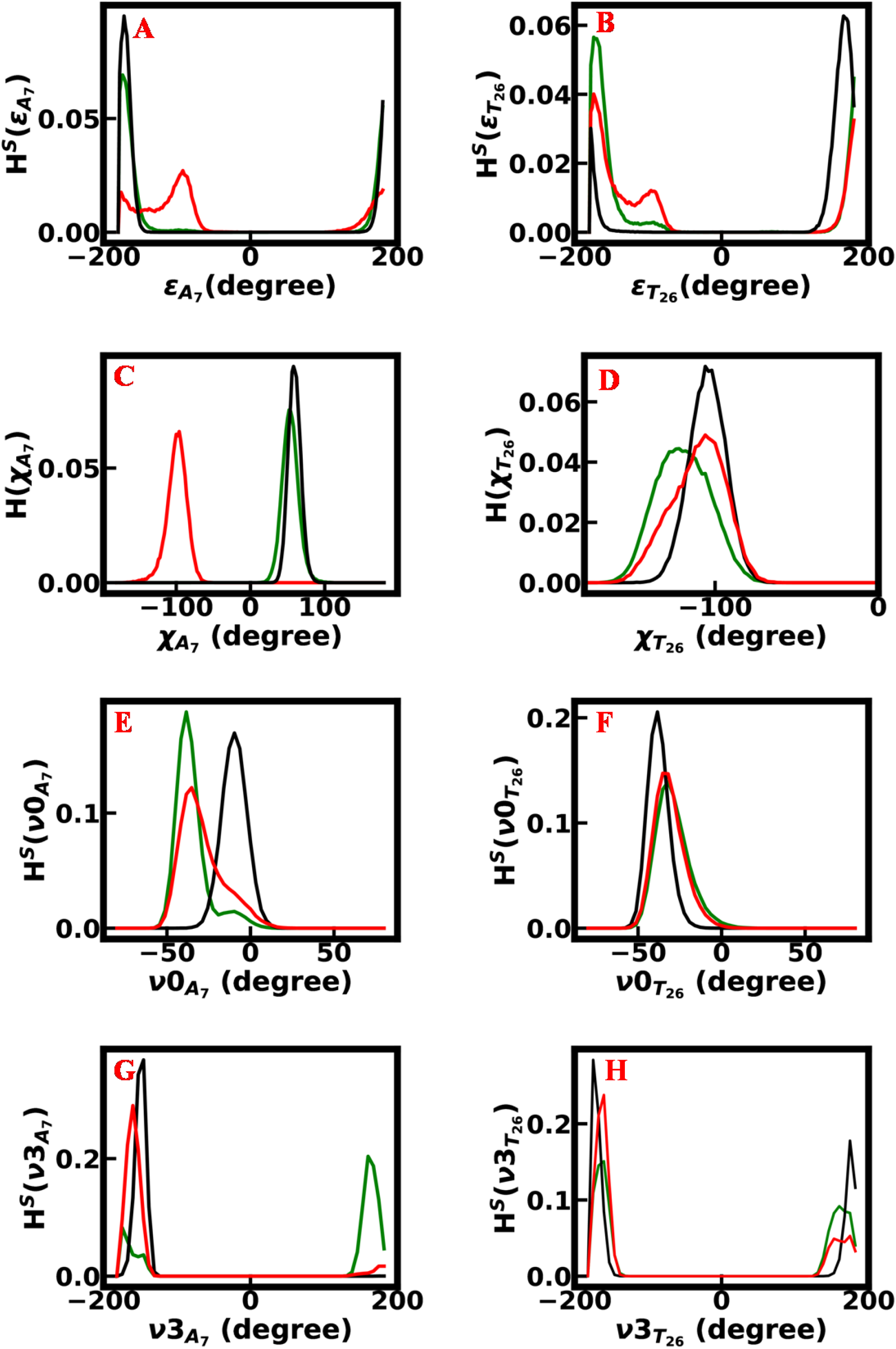
Histograms of torsion angles (A) ε for A_7_, (B) ε for T_26_, (C) χ for A_7_, (D) χ for T_26_ (E) ν0 of base A_7_, (F) ν0 of base T_26_, (G) ν3 for base A_7_, and (H) ν3 for base T_26_, for three systems in equilibrated trajectories. Each system is represented by a distinct color: the WC system is in red, the HG system is in green, and the HGP system is in black (S=WC, HG, and HGP).

Now we consider the distributions of the sugar torsion angles ν0 and ν3. Let us consider A_7_..In Fig. 2 (E), 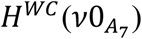 shows a single peak. The single peak changes to double peaks in 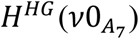, and reverts back to a single peak in 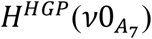. But, 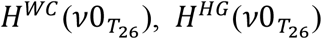 and 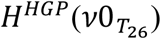 are single peaked (Fig. 2 (F)), the peak being the highest in 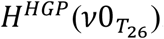. Again, we see, at A_7_, 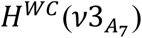 has one peak, but 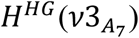 has two peaks, as illustrated in Fig. 2(G). However, 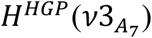 has a sharp peak higher than 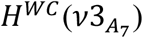 for A_7_. 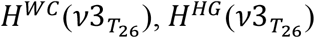 and 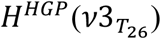, are all double peaked, each with a sharp peak (Fig. 2 (H)).The peak in 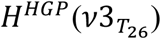 is the highest. Overall, the HG system shows multi-peaked or broad distribution in most of the microscopic conformational degrees of freedom compared to the WC system for A_7_-T_26_ bp. This suggests that HG bp is more flexible than WC bp in naked DNA. On the other hand, in the majority of cases, the sharper peaks of the HGP complex indicate that the presence of proteins reduce the flexibility of the HG bp.

### 3.3 Conformational thermodynamics

The changes in the distribution of the microscopic conformation variables lead to the changes in conformational free energy and entropy of DNA in HG and HGP systems with reference to the WC system. By adding the changes in the individual microscopic degrees of freedom, we compute the overall changes in conformational thermodynamics for the complete DNA duplex.

We first consider the changes in conformational thermodynamics in the inter-bp step parameters. For a given step parameter, Θ we denote the conformational free energy and entropy changes in the HG system with respect to the WC system by 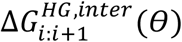 and 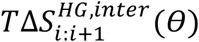 for the step i:i+1, where i and i+1 are the bases in 5’-3’ direction along with their complementary bases in the opposite strand. Figs. 3(A) and (B) show the data. We find that 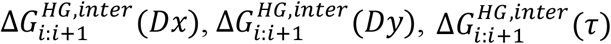 and 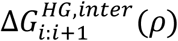 do not depend significantly on the bp steps in Fig. 3(A). On the other hand, 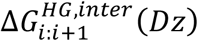 has maximum destabilization at the step A_7_:A_8_. 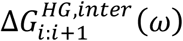 also contributes the most to destabilizing the step A_7_:A_8_. Let us now consider the changes in entropy. In Fig. 3(B), we see that 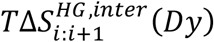 and 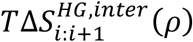 are not sensitive to bp steps. The step T_6_:A_7_ show the maximum order by 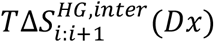, whereas the step A_7_:A_8_ is most disordered by 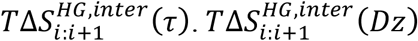 shows maximum disorder at the step T_6_:A_7_. However, 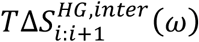 shows maximum order at the step T_6_:A_7_ and maximum disorder at the step A_7_:A_8_. We observe that due to inter-bp step parameters, the total changes in conformational free energy for a specific step, 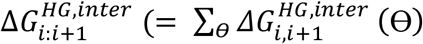, where Θ includes all of the step parameters) exhibits maximum destabilization at A_7_:A_8_ and total changes in conformation entropy, 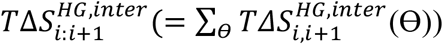 shows maximum conformational order at step T_6_:A_7_ and maximum conformational disorder at step A_7_:A_8_.

**Fig. 3:**
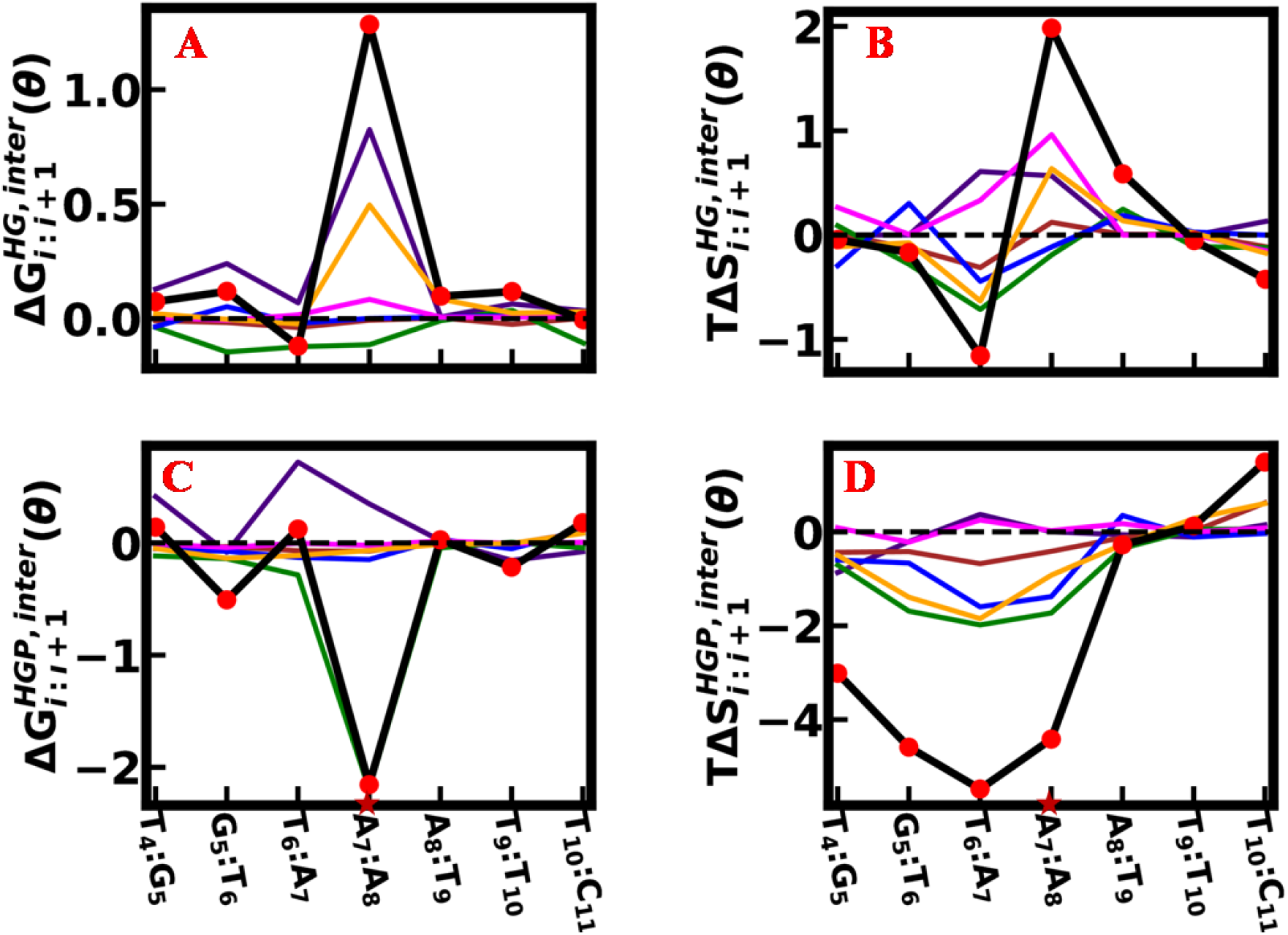
(A) 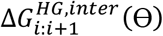 and (B) 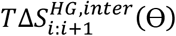, (C) 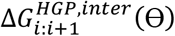 and (D) 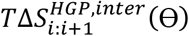, due to inter-bp step parameters for HG and HGP systems, respectively, with respect to the WC system. The distinct colors are used for different step parameters: D_x_ in green, D_y_ in blue, D_z_ in indigo, τ in magenta, ρ in brown and ω in orange. The total changes for each step are represented by the red circles connected by a solid black line. HG bp is marked by the maroon star symbol. All quantities are in kJ/mol.

The conformational free energy and entropy changes of the HGP system in comparison to the WC system for i:i+1 step are denoted by 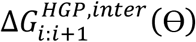 and 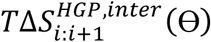 as shown in Figs. 3(C) and (D), respectively. Fig. 3(C) shows that 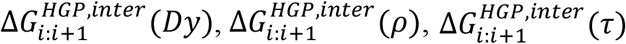 and 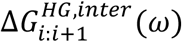 are not sensitive to the bp steps. 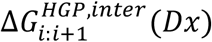 contributes the most to stabilizing the step A_7_:A_8_. Again, 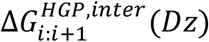 has maximum destabilization at the step T_6_:A_7_. Fig. 3(D) shows that 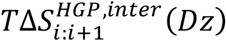 and 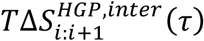 do not contribute significantly to the bp steps. 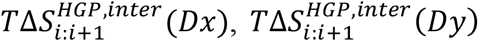 and 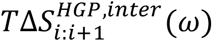 have maximum contributions to order the step G_5_:T_6_ to A_7_:A_8_. Again, 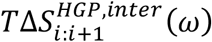 and 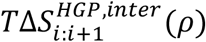 contribute maximum to disorder in the step T_10_:C_11_. The total changes in conformational free energy for every step, 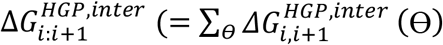, Θ running over all bp step parameters) exhibit maximum stabilization at the step A_7_:A_8_. The total changes in conformation entropy for individual step, 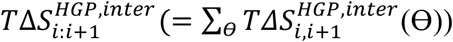 are shown in Fig. 3(D). The data show that there is significant ordering in the bp steps T_4_:G_5_ to A_7_:A_8_, with the maximum ordering at T_6_:A_7_.

The data for the changes in the conformational free energy and entropy, denoted by 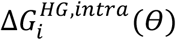 and 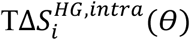, respectively, for the intra-bp parameter *θ* for *i*^*th*^ bp of the HG system with respect to the WC system are shown in Figs. 4(A) and (B), respectively.

**Fig. 4:**
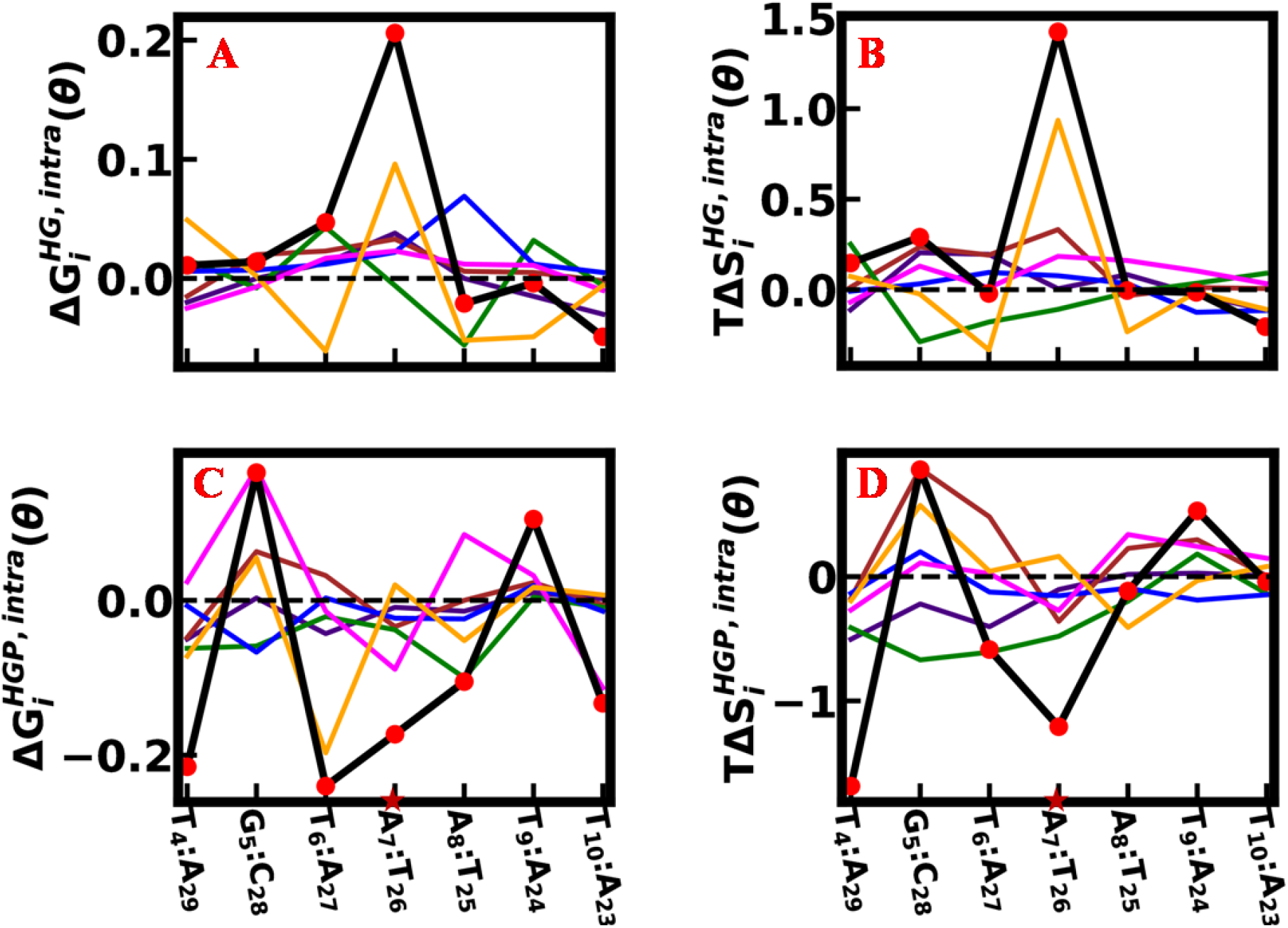
(A) 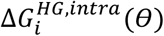 and (B) 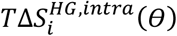, (C) 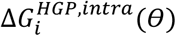 and (D) 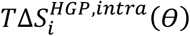, by intra-bp parameters, for HG and HGP systems, respectively, with respect to WC system. The distinct colors are used for different intra-bp parameters: S_x_ in magenta, S_y_ in blue, S_z_ in orange and κ in indigo, σ in brown, π in green. The total changes for each bp are represented by the red circles connected by a solid black line. HG bp is marked by the maroon star symbol. All quantities are in kJ/mol.

Here, 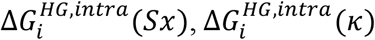 and 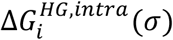 are not sensitive to the bp (Fig. 4(A)). However, 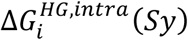 has maximum destabilization at A_8_:T_25_. Again, 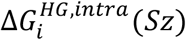 exhibits maximum destabilization at A_7_:T_26_ and maximum stabilization for T_6_:A_27_. However, 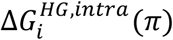 undergoes the maximum stabilization at A_8_:T_25_. Again, 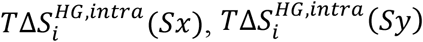 and 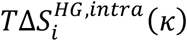 do not depend significantly on the bps (Fig.4(B)). However, 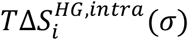 has maximum disorder at A_7_:T_26_. 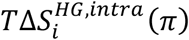 exhibits maximum order at G_5_:C_28_. Again, 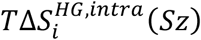 shows maximum disorder at A_7_:T_26_ and maximum order at T_6_:A_27_. Finally, we compute the total changes in conformational free energy, 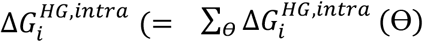, Θ running over the intra-bp parameters) and conformational entropy, 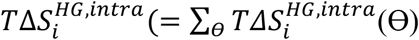, Θ runs over the intra-bp parameters) for each bp for intra-bp parameters. At A_7_:T_26_, 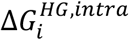 exhibits the most conformational destabilization, whereas 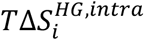 exhibits the most conformational disorder.

The changes in the conformational free energy and entropy for the HGP system with respect to the WC system, denoted by 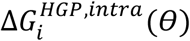 and 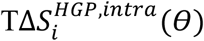, respectively, are shown in Figs. 4(C) and (D), respectively. In Fig. 4(C), we see that 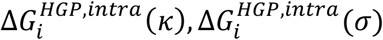 and 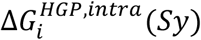 do not depend significantly to the bps. 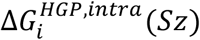 and 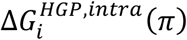 contribute maximum to stabilize the T_6_:A_27_ and A_8_:T_25_, respectively. Again, 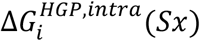 has maximum destabilization and stabilization for G_5_:C_28_ and T_10_:A_23_, respectively. In Fig. 4(D), we see that 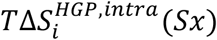 and 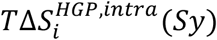 do not contribute significantly on the bps. 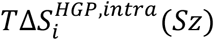 has maximum disorder at G_5_:C_28_ and maximum order at A_8_:T_25_. However, 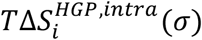 contributes maximum disorder to the step G_5_:C_28_. 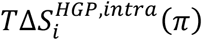 shows maximum order at G_5_:C_28_. Here, the total changes in conformational free energy for intra-bp parameters, 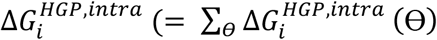,where, Θ runs over the intra-bp parameters) exhibit maximum conformational stabilization at T_6_:A_27_ and the total changes in conformational entropy for a specific bp, 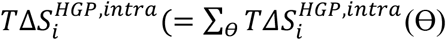, Θ runs over the intra-bp parameters) exhibit maximum order at T_4_:A_29_.

Next, we consider the conformational thermodynamic changes caused by the sugar-phosphate and sugar-base torsion angles. Purines and pyrimidines bases are present in the two strands of DNA. Again, in the HGP system, the interacting patterns of two proteins with strands are different. So, we compute strand specific conformational thermodynamics due to sugar-phosphate, sugar-base and sugar-puckers. The conformational free energy, 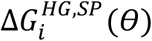 and entropy, 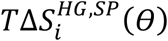 changes data for sugar-phosphate and sugar-base torsions for the *i*^*th*^ base of 5’-3’ strand of the HG system with respect to the WC system are shown in Figs. 5(A) and (B), respectively, where *θ* indicates sugar-base torsion angle χ along with sugar-phosphate backbone torsion angles α, β, δ, ε, γ and ζ. We find that 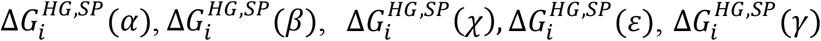 and 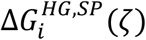 do not contribute significantly to bases in Fig. 5(A). 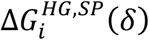 exhibits maximum destabilization and stabilization at A_7_ and G_5_, respectively. In Fig. 5(B), we see that 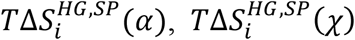 and 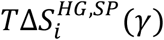 are not sensitive to the bases. However, 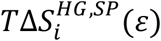 contributes the most to ordering the A_7_. On the other hand, 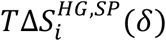 has maximum disorder at A_7_ while 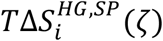 and 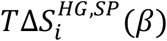 have maximum order at A_7_ and A_8_, respectively. We compute the total changes in conformational free energy, 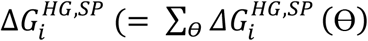, *θ* stands for sugar-base torsion angle χ along with sugar-phosphate backbone torsion angles α, β, δ, ε, γ and ζ) and conformational entropy, 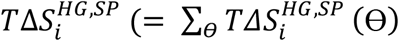, Θ runs over sugar-base torsion angle χ along with sugar-phosphate backbone torsion angles α, β, δ, ε, γ and ζ). 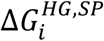 shows maximum destabilization at A_7_ while 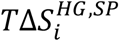 shows maximum order at A_7_.

**Fig. 5:**
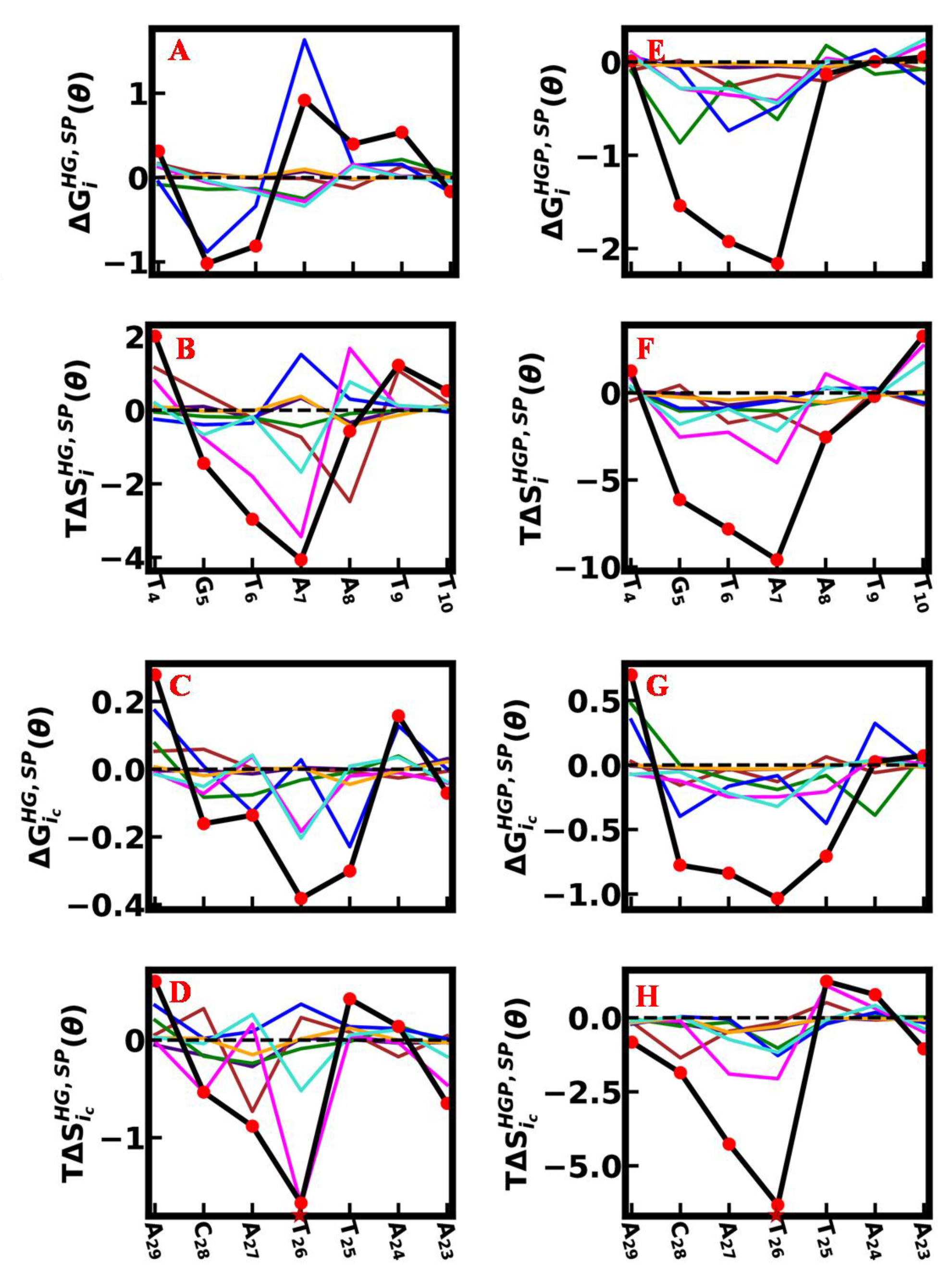
For sugar-phosphate and sugar-base backbone torsion angles, (A) 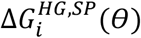 and (B) 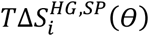 for 5’-3’ strand, (C) 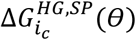 and (D) 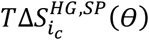 for 3’-5’ strand, in HG system with respect to WC system. Similarly, (E) 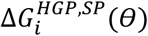 and (F) 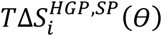 for 5’-3’ strand, (G) 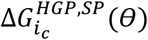 and (H) 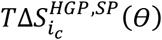 for 3’-5’ strand, in HGP system with regard to WC system. The distinct colors are used for different parameters for each Fig. : α in indigo, β in brown, χ in green, δ in blue, ε in magenta, γ in orange and ζ in turquoise. The total changes for each base are represented by the red circles connected by a solid black line. HG bp region is marked by the maroon star symbol. All quantities are in kJ/mol.

For the 3’-5’ strand, the conformational free energy, 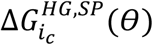 and entropy, 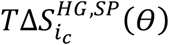 changes data for sugar-phosphate and sugar-base torsions for the *i*_*c*_^*th*^ base of the HG system with respect to the WC system are shown in Figs. 5(C) and (D), respectively. In Fig. 5(C), 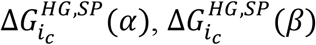 and 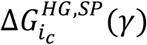 are not sensitive to bases. In contrast, 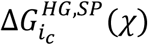 has maximum stability at C_28_ while 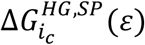 and 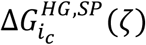 have maximum stability at T_26_. 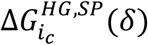 shows maximum destabilization and stabilization at A_29_ and T_25_, respectively. On the other hand, in Fig. 5(D), 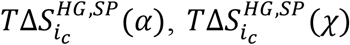 and 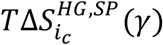 are not sensitive to bases. 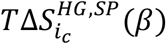 has maximum order at A_27_. 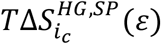 and 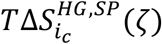 show maximum order at T_26_ while 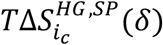 shows maximum disorder at T_26_. We evaluate the total changes in conformational free energy, 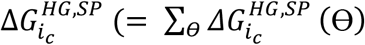, *θ* stands for sugar-base torsion angle χ along with sugar-phosphate backbone torsion angles α, β, δ, ε, γ and ζ) and conformational entropy, 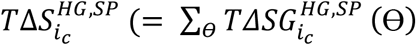, *θ* runs over sugar-base torsion angle χ along with sugar-phosphate backbone torsion angles α, β, δ, ε, γ and ζ) for 3’-5’ strand. Here, 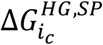 shows maximum stabilization at T_26_ while 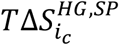 shows maximum order at T_26_.

For sugar-phosphate and sugar-base torsion angles, the conformational free energy and entropy, 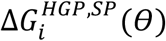, and 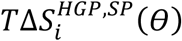 for i^th^ base along 5’-3’ strand in the HGP system with regard to the WC system are shown in Figs. 5(E) and (F), respectively. In Fig. 5(E), we observe that 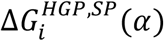, 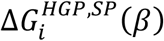 and 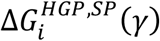 are not sensitive to the bases. 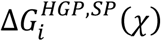 contributes the most to stabilize the G_5_. 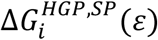 and 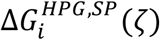 show maximum stabilization at A_7_. Again, 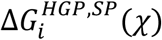 contributes the most to stabilizing the A_7_. However, 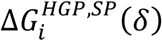 has maximum contribution to stabilizing T_6_. We find in Fig. 5(F) that 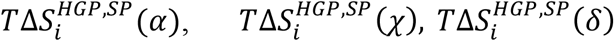 and 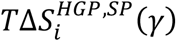 do not contribute significantly to the bases. Otherwise, 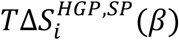 has maximum order at A_8_. 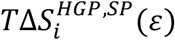 contributes maximum to order the A_7_, here. Again, 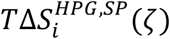 has maximum order and disorder at A_7_ and T_10_, respectively. The total changes in conformational free energy, 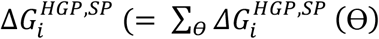, *θ* runs over sugar-base torsion angle χ along with sugar-phosphate backbone torsion angles α, β, δ, ε, γ and ζ) shows stabilization from G_5_ to A_7_ with a maximum value at A_7_ and the total changes in conformational entropy, 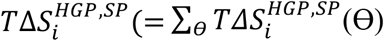, *θ* runs over sugar-base torsion angle χ along with sugar-phosphate backbone torsion angles α, β, δ, ε, γ and ζ) shows strong ordering from G_5_ to A_7_, with a maximum value at A_7_.

In the complementary 3’-5’ strand, the conformational free energy and entropy changes for the HGP system with respect to the WC system caused by sugar-phosphate and sugar-base torsion angles, 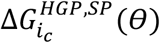 and 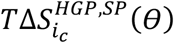 for the base 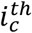 are shown in Figs. 5(G) and (H), respectively. 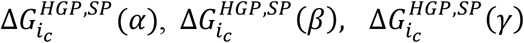 and 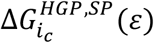 do not show sensitivity to the bases in Fig. 5(G). On the other hand, 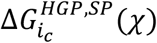 has maximum destabilization and stabilization at A_29_ and A_24_, respectively. Again, 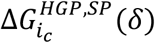 shows maximum destabilization and stabilization at A_29_ and T_25_, respectively. At T_26_, 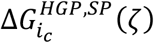 shows maximum stabilization. In Fig. 5(H), 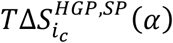 and 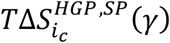 are not sensitive to bases. However, 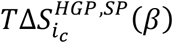 contributes the most to ordering the C_28_. On the other hand, 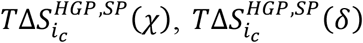 and 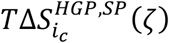 contribute maximum to order the T_26_. Again, 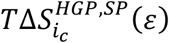 contributes the most to organizing the T_26_. Thus, for the complementary strand, the total changes in conformational free energy, 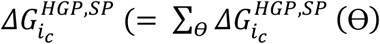, *θ* runs over sugar-base torsion angle χ along with sugar-phosphate backbone torsion angles α, β, δ, ε, γ and ζ) demonstrate the largest stability at T_26_. On the other hand, the total changes in conformational entropy, 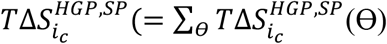, *θ* runs over sugar-base torsion angle χ along with sugar-phosphate backbone torsion angles α, β, δ, ε, γ and ζ) show significant ordering from C_28_ to T_26_, with maximum value at T_26_.

In the 5’-3’ strand, the changes in conformational free energy, 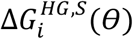 and entropy, 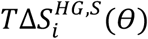 due to the sugar torsion angles (ν0, ν1, ν2, ν3 and ν4) of each of *i*^*th*^ base in the HG system with respect to the WC system are shown in Figs. 6(A) and (B), respectively. We note in Fig. 6(A) that 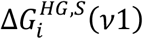 is not sensitive to bases. At G_5_ and A_7_, 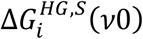 exhibits the most destabilization and stabilization, respectively. On the other hand, 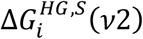 contributes the most to destabilizing the A_7_ and to stabilizing the T_6_. However, 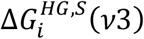 has maximum destabilization for A_7_. Again, 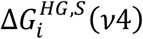 contributes maximum to destabilize A_8_. We find that in Fig. 6(B), 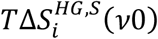 and 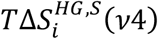 have maximum order to A_7_. 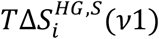 contributes the most to disorder the A_8_. 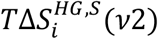 shows maximum disorder at A_7_. However, 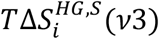 contributes the most to order the T_6_ and to disorder the A_7_. We compute the total changes in conformational free energy, 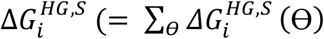, Θ runs over all sugar-puckers) and conformational entropy, 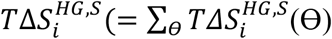, Θ runs over all sugar-puckers). 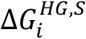 exhibits maximum stabilization at T_6_ while 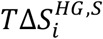 shows maximum disorder at A_8_.

**Fig. 6:**
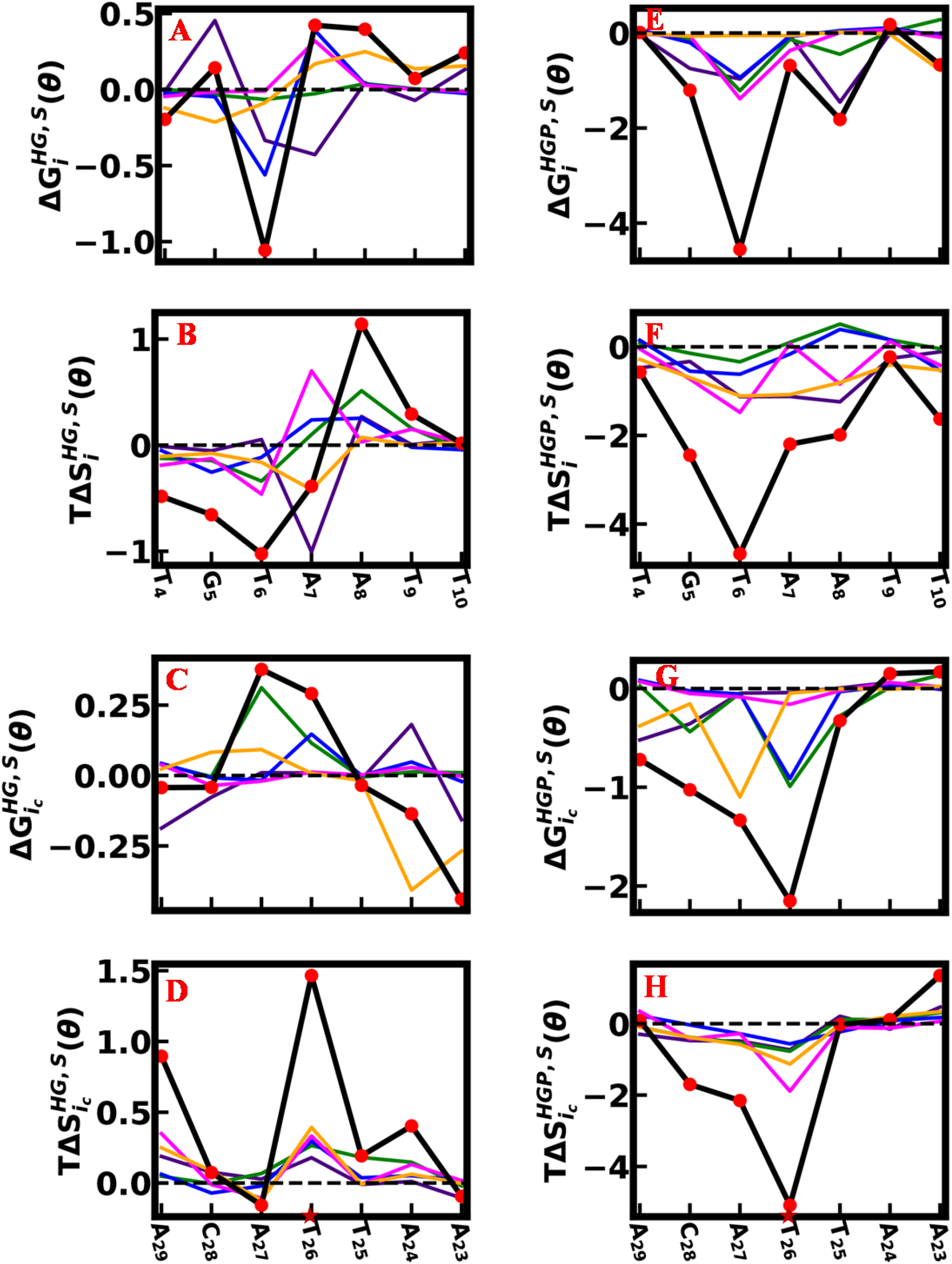
For sugar-puckers, (A) 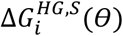 and (B) 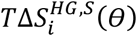 for 5’-3’ strand, (C) 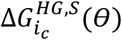 and (D) 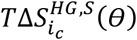 for 3’-5’ strand, in HG system with respect to WC system. Similarly, (E) 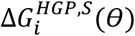 and (F) 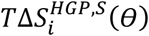 for 5’-3’ strand, (G) 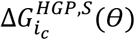 and (H) 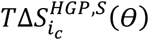 for 3’-5’ strand, in HGP system with regard to WC system. The distinct colors are used for different parameters in each Fig. : ν0 in indigo, ν1 in green, ν2 in blue, ν3 in magenta and ν4 in orange. The total changes for each base are represented by the red circles connected by a solid black line. HG bp region is marked by the maroon star symbol. All quantities are in kJ/mol.

In the 3’-5’ strand, the changes in conformational free energy, 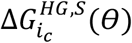 and entropy, 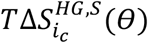 due to the sugar torsion angles of each of *i*_*c*_^*th*^ base in the HG system with regard to the WC system are shown in Figs. 6(C) and (D), respectively. According to Fig. 6(C), 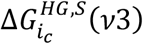 is not sensitive to bases. 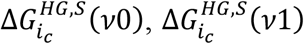 and 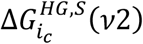 show maximum destabilization at A_24_, A_27_ and T_26_, respectively. Again, 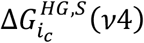 shows maximum stabilization at A_24_. In Fig. 6(D), 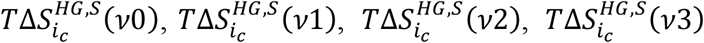 and 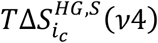 show maximum disorder at T_26_. We calculate the total changes in conformational free energy, 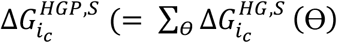, Θ runs over all sugar-puckers) and conformational entropy, 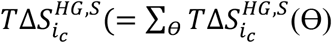, Θ runs over all sugar-puckers) for the complementary 3’-5’ strand. 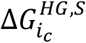 shows maximum stabilization at A_23_ while 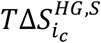 shows maximum disorder at T_26_.

In the HGP system with regard to the WC system, the changes in conformational free energy, 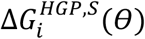 and entropy, 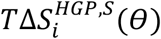 due to the sugar torsion angles, of *i*^*th*^ base of 5’-3’ strand are shown in Figs. 6(E) and (F), respectively. In Fig. 6(E), 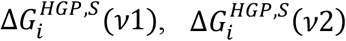 and 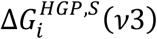 exhibit maximum stabilization at T_6_. 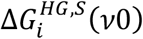 has maximum contribution to stabilize the G_5_ and A_8_. 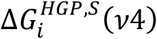 has maximum destabilization at T_10_. We find that in Fig. 6(F), 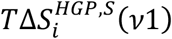 and 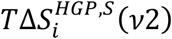 do not sensitive to bases. But, 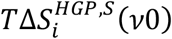 and 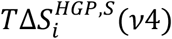 contribute the most to ordering the A_7_. Again, 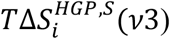 has maximum contribution to order the T_6_. The total changes in conformational free energy, 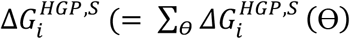, Θ runs over all sugar-puckers) exhibit significant stabilization at T_6_ and A_8_, while the total changes in conformational entropy, 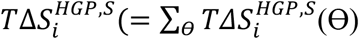, Θ runs over all sugar-puckers) show maximum order at T_6_.

For sugar pucker angles, the changes in conformational free energy, 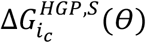 and entropy, 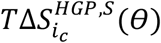 of *i*_*c*_^*th*^ base of the complementary 3’-5’ strand of the HGP system with regard to the WC system are shown in Figs. 6(G) and (H), respectively. In Fig, 6(G), 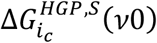 and 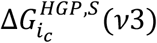 do not contribute significantly to bases. However, 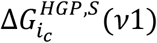 and 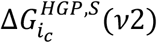 show maximum stabilization at T_26_. Again, 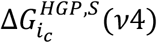 exhibits the highest stability at A_27_. In Fig. 6(H), 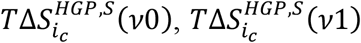 and 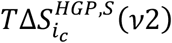 are not sensitive to the bases. Only, 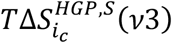 and 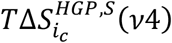 have maximum ordering at T_26_. Here, the total changes in conformational free energy, 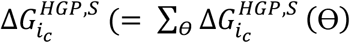, Θ runs over all sugar-puckers) show the most stabilization at T_26_ while the total changes in conformational entropy, 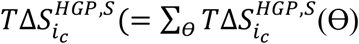, Θ runs over all sugar-puckers) exhibit significant ordering from C_28_ to T_26_, with a maximum value at T_26_.

We compute the changes in conformational free energy and entropy, 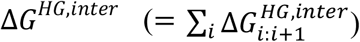 and 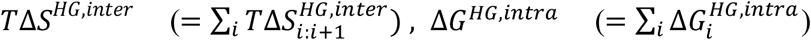 and 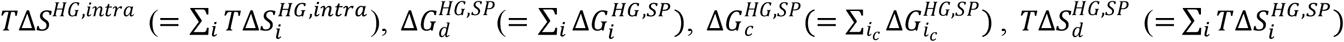 and 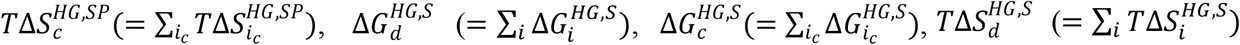 and 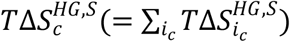 for inter-bp step parameters, intra-bp parameters, sugar-base, sugar-phosphate torsion angles, and sugar-puckers, respectively, in the HG system with regard to the WC system. Here, “d” refers to the 5’-3’ strand and “c” to the complementary 3’-5’ strand. The data are shown in Table 1. We find that most of the changes in conformational free energy and entropy are insignificant with respect to thermal energy (≈2.5 kJ/mol). Only the conformational entropy, 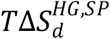 and 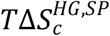 show more ordering due to sugar-base and sugar-phosphate torsion angles.

**Table 1:** The changes (kJ/mol) in conformational thermodynamics of HG and HGP systems in comparison to the WC system.

| $\Delta G^{HG,inter}$ | $T\Delta S^{HG,inter}$ | $\Delta G^{HG,intra}$ | $T\Delta S^{HG,intra}$ | $\Delta G_d^{HG,SP}$ | $\Delta G_c^{HG,SP}$ | $T\Delta S_d^{HG,SP}$ | $T\Delta S_c^{HG,SP}$ | $\Delta G_d^{HG,S}$ | $\Delta G_c^{HG,S}$ | $T\Delta S_d^{HG,S}$ | $T\Delta S_c^{HG,S}$ | $\Delta G_{total}^{HG}$ | $T\Delta S_{total}^{HG}$ |
| --- | --- | --- | --- | --- | --- | --- | --- | --- | --- | --- | --- | --- | --- |
| 1.57 | 0.73 | 0.20 | 1.62 | 0.17 | -0.61 | -5.27 | -2.57 | 0.02 | -0.03 | -1.1 | 2.78 | 1.33 | -3.81 |
| $\Delta G^{HGP,inter}$ | $T\Delta S^{HGP,inter}$ | $\Delta G^{HGP,intra}$ | $T\Delta S^{HGP,intra}$ | $\Delta G_d^{HGP,SP}$ | $\Delta G_c^{HGP,SP}$ | $T\Delta S_d^{HGP,SP}$ | $T\Delta S_c^{HGP,SP}$ | $\Delta G_d^{HGP,S}$ | $\Delta G_c^{HGP,S}$ | $T\Delta S_d^{HGP,S}$ | $T\Delta S_c^{HGP,S}$ | $\Delta G_{total}^{HGP}$ | $T\Delta S_{total}^{HGP}$ |
| -2.41 | -16.16 | -0.60 | -2.25 | -5.69 | -2.56 | -21.74 | -12.31 | -8.74 | -5.24 | -13.74 | -7.42 | -25.23 | -73.62 |

Similarly, we calculate 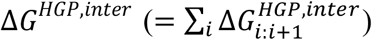 and 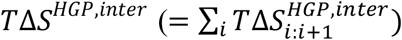 for inter-bp parameters, 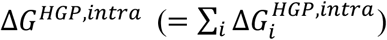 and 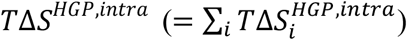 for intra-bp parameters, 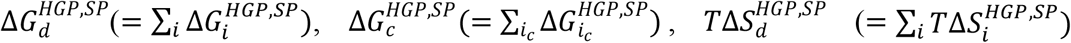 and 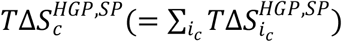 for sugar-base and sugar-phosphate backbone torsion angles, 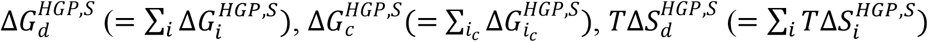 and 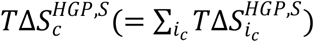 due to sugar-puckers, in the HGP system with respect to the WC system. Here, “d” stands to the 5’-3’ strand and “c” to the complementary 3’-5’ strand. Table 1 shows stabilization more in 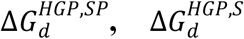 and 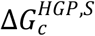 and ordering significantly in 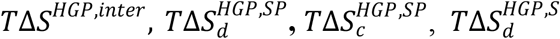 and 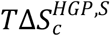, in the HGP system with respect to the WC system.

Finally, we compute 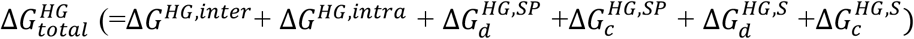 and 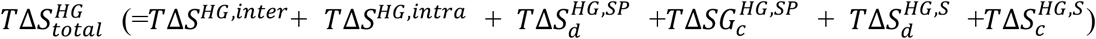 values, for the HG system with respect to the WC system are 1.33 kJ/mol and -3.81 kJ/mol, respectively (Table 1), while values of 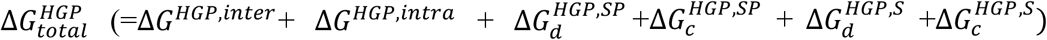 and 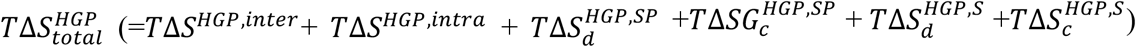 for the HGP system in comparison to the WC system are -25.23 kJ/mol and -73.62 kJ/mol, respectively (Table 1). Overall, the total changes of conformational thermodynamics of the entire DNA system, we observe that the HG system is slightly destabilized but ordered than the WC system, while the HGP complex is significantly stabilized and ordered in comparison to the WC system.

The inter-bp step parameters are the primary factors to destabilize the HG system from the WC system, while sugar-puckers are the main factors to stabilize the HGP system in contrast to the WC system. From Table 1, we observe that for the 5’-3’ strand, 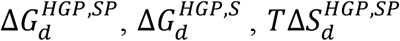 and 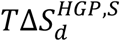 are more negative compared to 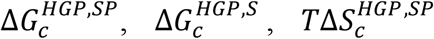 and 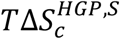, respectively, of the complementary 3’-5’ strand. This shows that protein α2D has a greater impact on stabilizing and organizing the HGP system than protein α2B. Based on the fluctuation of the dihedral angle χ alone in the presence and absence of proteins, the earlier MD simulation on this system reveals that the non-specifically bound protein α2D has a stronger influence on the formation of HG bp^11^. In addition to χ, here we observe that the HG bp in the HGP system stabilized and ordered more for the microscopic degrees of freedom D_x_, δ, ε, ζ, ν0, ν1, ν2, ν3 and ν4. This gives a more complete microscopic picture of the stability at the HG bp induced by the bound protein.

## 4. Conclusions

To summarize, the fluctuations of microscopic conformational variables reveal that the HG bp forming region as well as the entire DNA duplex become stabilized and ordered in the presence of a non-specifically bound protein α2D and a specifically bound protein α2B in the 1K61 system. Protein α2D is more effective than protein α2B at stabilizing and organizing the HGP system. Such study based on the probability distribution of proper conformational variable is a new approach to understanding the stability of such variable bp geometry due to non-canonical HG base pairing. Furthermore, the modulation of DNA-protein recognition due to non-canonical base pairing can also be implemented on the stability and order of tumor suppressor proteins like p53 or DNA lesion repair proteins interfered with by HG base pairing. It is highly interesting to analyze their effect on signal transduction and mismatch repair systems effectively in the future. Our study can also lead to a focus on anti-cancer drug target therapeutics acting through DNA mediated HG base pairing.

## 6. Acknowledgement

KK thanks to CSIR-UGC for providing the fellowship to conduct the research and Dr. Sasthi Charan Mandal for some helpful discussion. AMG is thankful to Council for Scientific and Industrial Research for financial support through Research Associateship (File No. 09/0575(11595)/2021-EMR-I).

## Supplementary information

### All the microscopic conformational variables of DNA

#### DNA step parameters

There are six DNA bp step parameters, three are rotational parameters (tilt (τ), roll (ρ), and twist (ω)) and the other three are translational parameters (shift (D_x_), slide (D_y_), and rise (D_z_))^1^. These are describing the position and orientation of one bp relative to the next bp. Shift is defined as the displacement of one bp with respect to the next along the X axis, while slide is the displacement about the Y axis. Rise is described as the displacement along the Z axis. Tilt, roll, and twist are defined as the rotation of the bp around shift, slide, and rise axes. The 5’-3’ direction of strand 1 is used as the positive Z direction. The long axis (Y-axis) can be along the C6-C8 direction and passes through the bp center. The bp center is defined as the midpoint of C6 and C8 atoms of pyrimidines and purines respectively. The short axis (X-axis) points towards the major groove. The DNA bp step parameters are calculated using the formula:

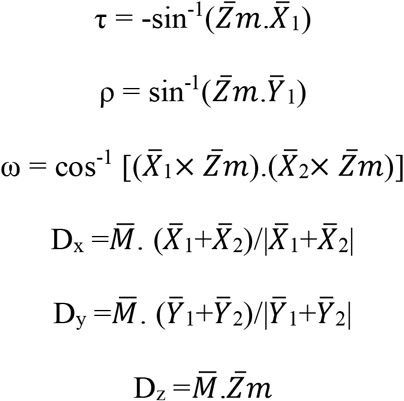

Here 1 and 2 denote two successive bp planes. 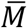 is the vector joining the bp centers of two consecutive bps. 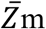 is defined in the following formula,

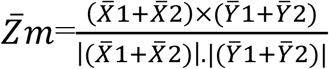

#### DNA intra-base pair parameters

There are six DNA base pair parameters; three are rotational parameters (buckle (κ), propeller (π), and opening (σ)) and three are translational parameters (stagger (S_x_), shear (S_y_), and stretch (S_z_))^2^. These are describing the position and orientation of one base relative to other bases in a bp. The axis systems for the two bases are in a pair following the hydrogen bonding edge of the bases involved^2^. The base’s X-axis is fixed perpendicular to the best mean plane across the base ring atoms, which can be along or against the 5’-3’ strand direction. Each base in a pair’s Y-axis is defined by two hydrogen-bonding heavy atoms of the specific edge forming the hydrogen bonds. A base’s Z-axis is set perpendicular to both the X- and Y-axes in a right-handed axis system, and it passes roughly parallel to the hydrogen bonds formed in the pair. The mathematical expression used in calculating buckle is analogous to that used for calculating tilt, while the expressions for opening, propeller, shear, stagger and stretch are identical to those used for calculating roll, twist, slide, shift and rise, respectively, but with the axes for the bases rather than the bps. The DNA bp parameters are calculated using the formula:

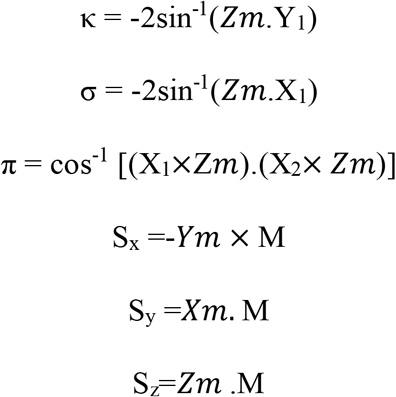

where X_1_, Y_1_ and Z_1_ are unit vectors along the axes of the first base, X_2_, Y_2_ and Z_2_ are those for the second base. The vector M is created by combining two base atoms, one from each base in the pair and chosen based on the base’s specific hydrogen-bonding edge. The components of the mean unit vector Xm, Ym and Zm are calculated as follows:

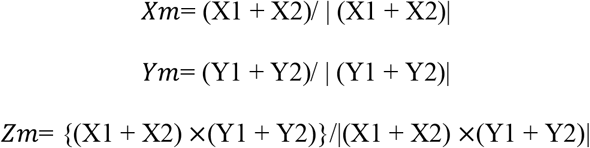

#### Sugar-phosphate and sugar-base torsion parameters

In DNA, there are six types of sugar-phosphates (alpha (α), beta (β), gamma (γ), delta (δ), epsilon (ε) and zeta (ζ)) and one type of sugar-base (chi (χ)) torsion angles^3^. They are as follows:

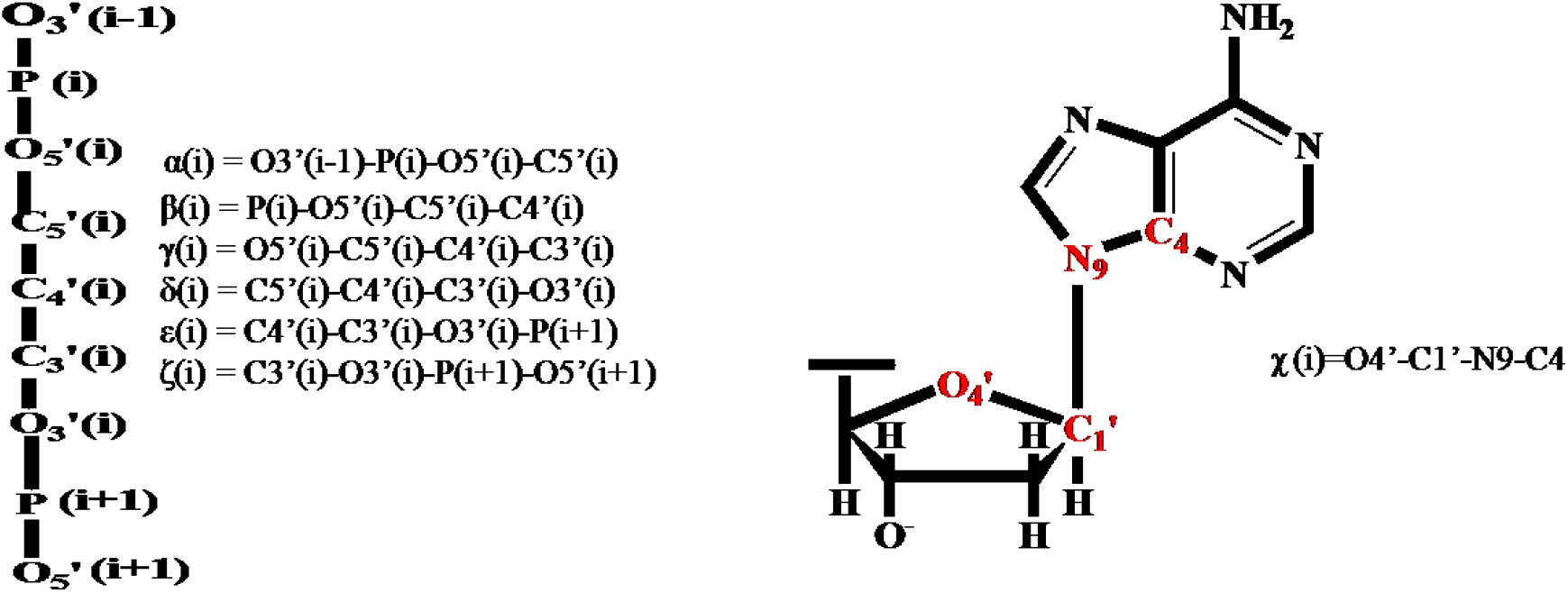

#### Sugar-pucker angles

For each sugar in DNA, there are five torsion angles (nu0 (ν0), nu1 (ν1), nu2 (ν2), nu3 (ν3) and nu4 (ν4)^4^. They are as follows:

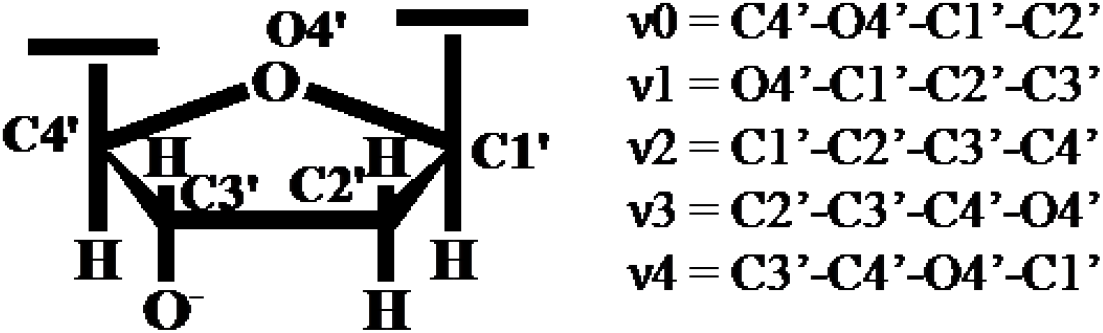

#### Pseudo rotation phase angle (P)

The five sugar pucker angles (ν0, ν1, ν2, ν3 and ν4) are used to determine the pseudo rotation phase angle (P) for a sugar^4^. It is defined as:

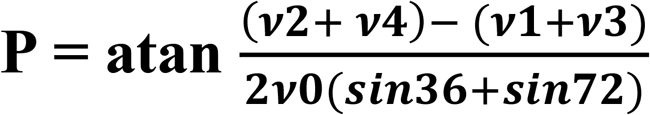

P is classified as C2’-endo when it falls between 137^0^ and 194^0^ and as C3’-exo when it falls between 195^0^ and 216^0^.

**Fig. S1:**
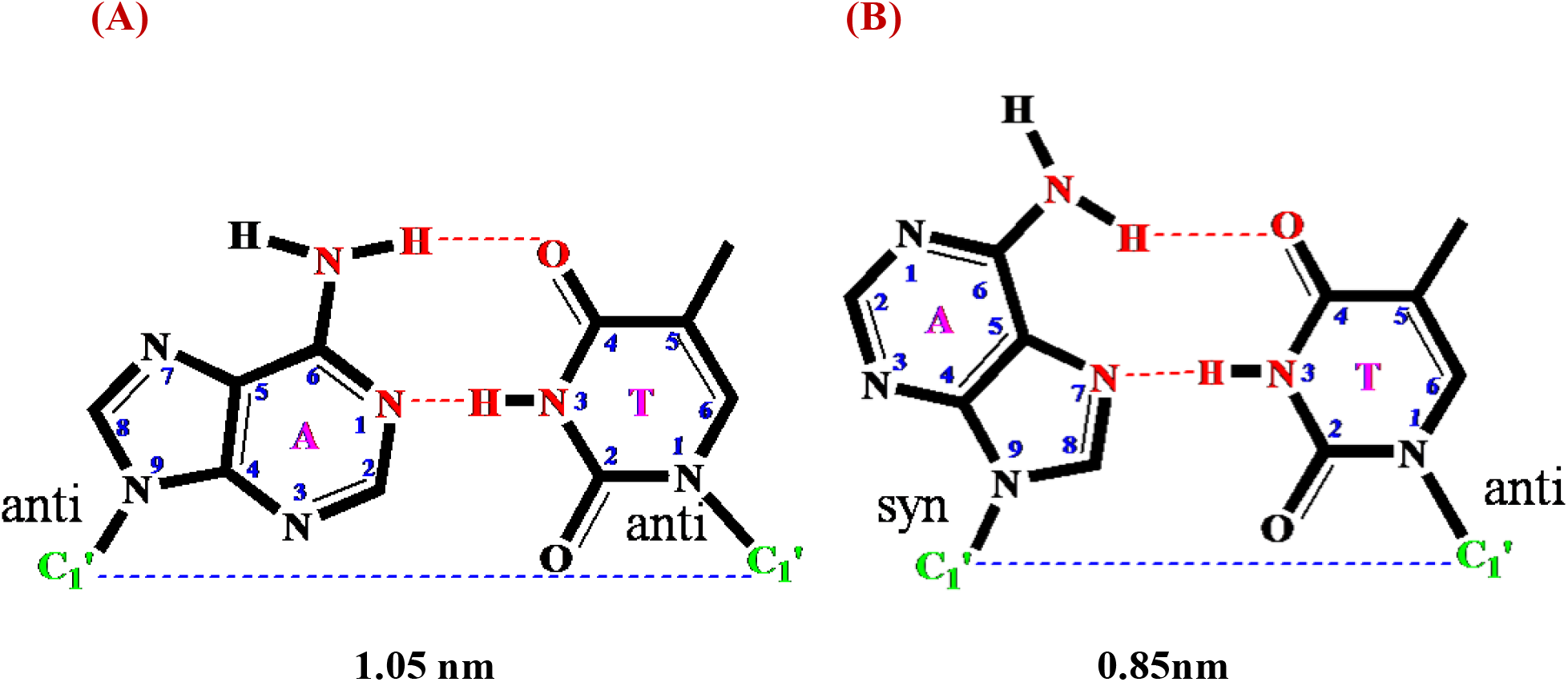
(A) WC and (B) HG A-T bp. Heavy atoms involved in hydrogen bonds are highlighted in red.

**Fig. S2:**
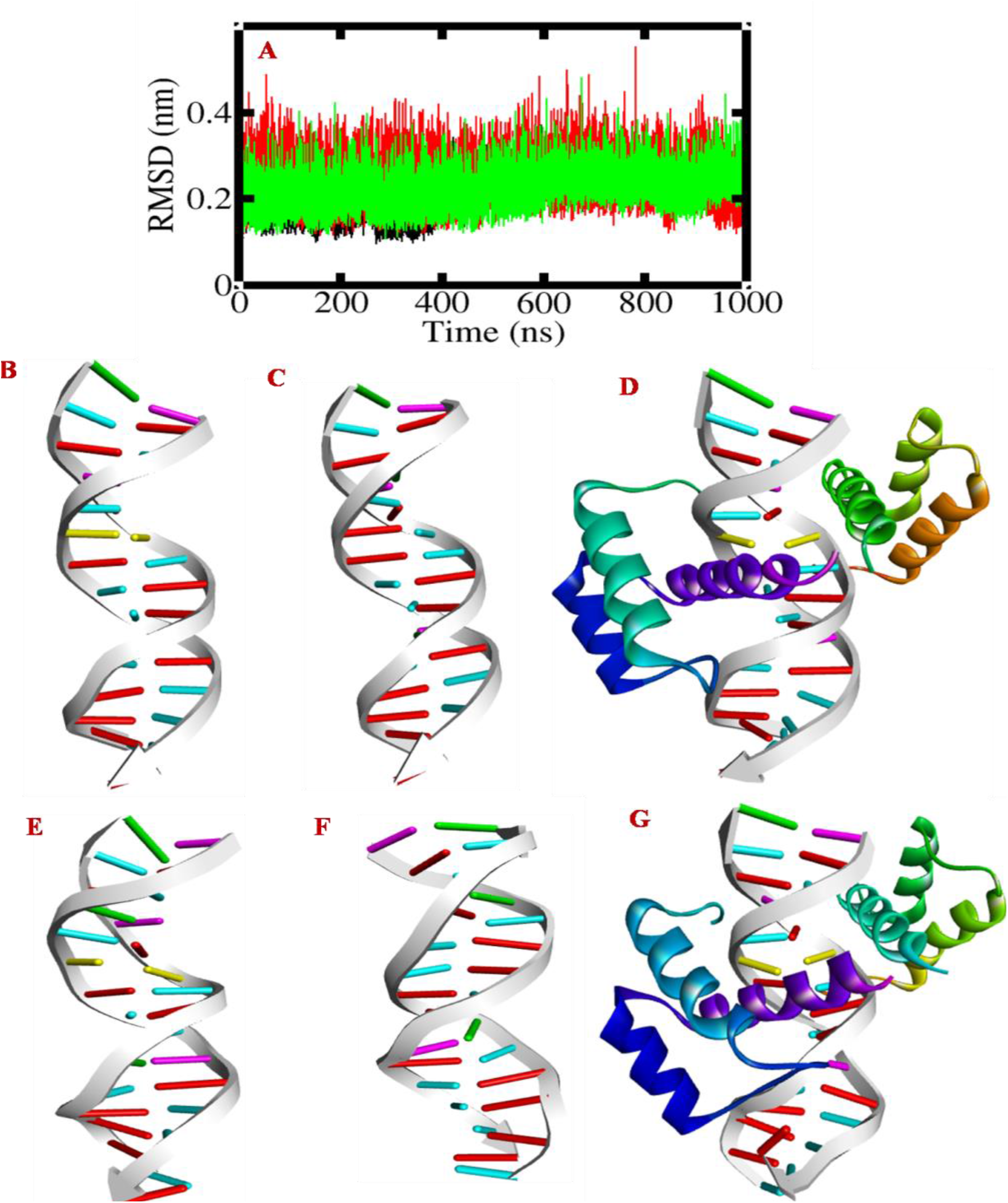
(A) RMSD of simulated ensembles, WC (red), HG (green), and HGP (black). Snapshot showing the initial structure of (B) HG (C) WC (D) HGP systems and snapshot at 1μs time span of (E) HG (F) WC (G) HGP systems. HG bp is highlighted in yellow.

**Fig. S3:**
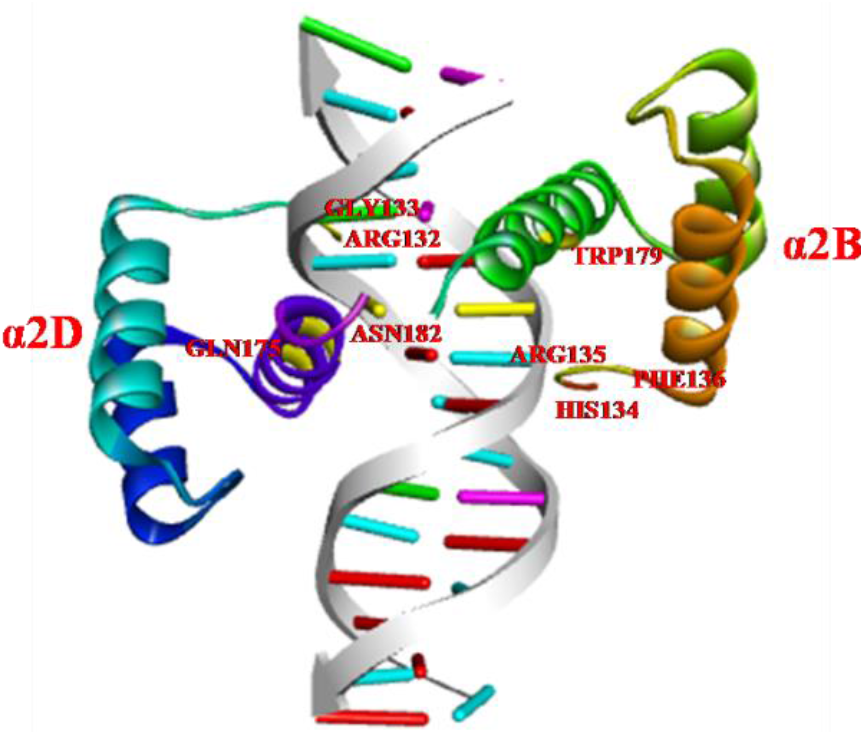
Snapshot of active protein residues at the interface in HGP system. Protein interface binding residues with HG bp are shown in yellow.

**Table S1:** The mean with the error in the mean of each step in the WC system for the bp step parameters (D_x_, D_y_, D_z_, τ, ρ, ω). D_x_, D_y_, D_z_ are in A^0^. τ, ρ and ω are in degree.

| System | steps | $(\langle D_x^{WC} \rangle \pm 0.5\Delta D_x^{WC})$ | $(\langle D_y^{WC} \rangle \pm 0.5\Delta D_y^{WC})$ | $(\langle D_z^{WC} \rangle \pm 0.5\Delta D_z^{WC})$ | $(\langle \tau^{WC} \rangle \pm 0.5\Delta \tau^{WC})$ | $(\langle \rho^{WC} \rangle \pm 0.5\Delta \rho^{WC})$ | $(\langle \omega^{WC} \rangle \pm 0.5\Delta \omega^{WC})$ |
| --- | --- | --- | --- | --- | --- | --- | --- |
| WC | T4-A29/G5-C28 | $0.40 \pm 0.008$ | $-0.17 \pm 0.01$ | $3.37 \pm 0.001$ | $0.47 \pm 0.04$ | $7.19 \pm 0.11$ | $37.68 \pm 0.04$ |
| | G5-C28/T6-A27 | $-0.43 \pm 0.01$ | $-0.19 \pm 0.003$ | $3.29 \pm 0.002$ | $0.11 \pm 0.02$ | $0.30 \pm 0.04$ | $31.35 \pm 0.05$ |
| | T6-A27/A7-T26 | $0.49 \pm 0.03$ | $0.24 \pm 0.01$ | $3.43 \pm 0.002$ | $1.82 \pm 0.08$ | $4.24 \pm 0.11$ | $32.65 \pm 0.12$ |
| | A7-T26/A8-T25 | $-0.62 \pm 0.01$ | $-0.02 \pm 0.01$ | $3.28 \pm 0.001$ | $-2.35 \pm 0.02$ | $0.59 \pm 0.11$ | $36.84 \pm 0.03$ |
| | A8-T25/T9-A24 | $0.02 \pm 0.003$ | $-0.69 \pm 0.002$ | $3.10 \pm 0.003$ | $0.02 \pm 0.01$ | $-0.73 \pm 0.10$ | $35.97 \pm 0.06$ |
| | T9-A24/T10-A23 | $0.32 \pm 0.005$ | $-0.23 \pm 0.01$ | $3.28 \pm 0.001$ | $1.80 \pm 0.02$ | $1.35 \pm 0.10$ | $37.34 \pm 0.04$ |
| | T10-A23/C11-G22 | $0.50 \pm 0.008$ | $0.17 \pm 0.01$ | $3.39 \pm 0.002$ | $0.78 \pm 0.02$ | $2.66 \pm 0.10$ | $32.45 \pm 0.10$ |

**Table S2:** The mean with the error in the mean of each step in the HG system for the bp step parameters (D_x_, D_y_, D_z_, τ, ρ, ω). D_x_, D_y_, D_z_ are in A^0^. τ, ρ and ω are in degree.

| System | steps | $\langle D_x^{HG} \rangle \pm 0.5\Delta D_x^{HG}$ | $\langle D_y^{HG} \rangle \pm 0.5\Delta D_y^{HG}$ | $\langle D_z^{HG} \rangle \pm 0.5\Delta D_z^{HG}$ | $\langle \tau^{HG} \rangle \pm 0.5\Delta \tau^{HG}$ | $\langle \rho^{HG} \rangle \pm 0.5\Delta \rho^{HG}$ | $\langle \omega^{HG} \rangle \pm 0.5\Delta \omega^{HG}$ |
| --- | --- | --- | --- | --- | --- | --- | --- |
| HG | T4-A29/G5-C28 | 0.06 ± 0.01 | -0.24 ± 0.01 | 3.33 ± 0.002 | 0.62 ± 0.03 | 7.41 ± 0.05 | 38.91 ± 0.05 |
| | G5-C28/T6-A27 | -0.23 ± 0.01 | -0.36 ± 0.01 | 3.19 ± 0.002 | 0.80 ± 0.01 | 0.35 ± 0.03 | 33.44 ± 0.05 |
| | T6-A27/A7-T26 | -0.48 ± 0.02 | -1.40 ± 0.01 | 3.30 ± 0.004 | -3.13 ± 0.04 | 2.75 ± 0.08 | 49.29 ± 0.08 |
| | A7-T26/A8-T25 | 1.20 ± 0.01 | 0.46 ± 0.01 | 3.61 ± 0.003 | 5.25 ± 0.06 | 3.75 ± 0.06 | 21.88 ± 0.08 |
| | A8-T25/T9-A24 | -0.04 ± 0.005 | -0.66 ± 0.004 | 3.15 ± 0.003 | -0.34 ± 0.01 | -1.61 ± 0.05 | 34.68 ± 0.08 |
| | T9-A24/T10-A23 | 0.20 ± 0.01 | -0.37 ± 0.01 | 3.28 ± 0.002 | 1.70 ± 0.02 | 1.46 ± 0.03 | 37.01 ± 0.03 |
| | T10-A23/C11-G22 | 0.56 ± 0.01 | 0.18 ± 0.01 | 3.38 ± 0.001 | 0.97 ± 0.02 | 2.57 ± 0.02 | 33.22 ± 0.04 |

**Table S3:**
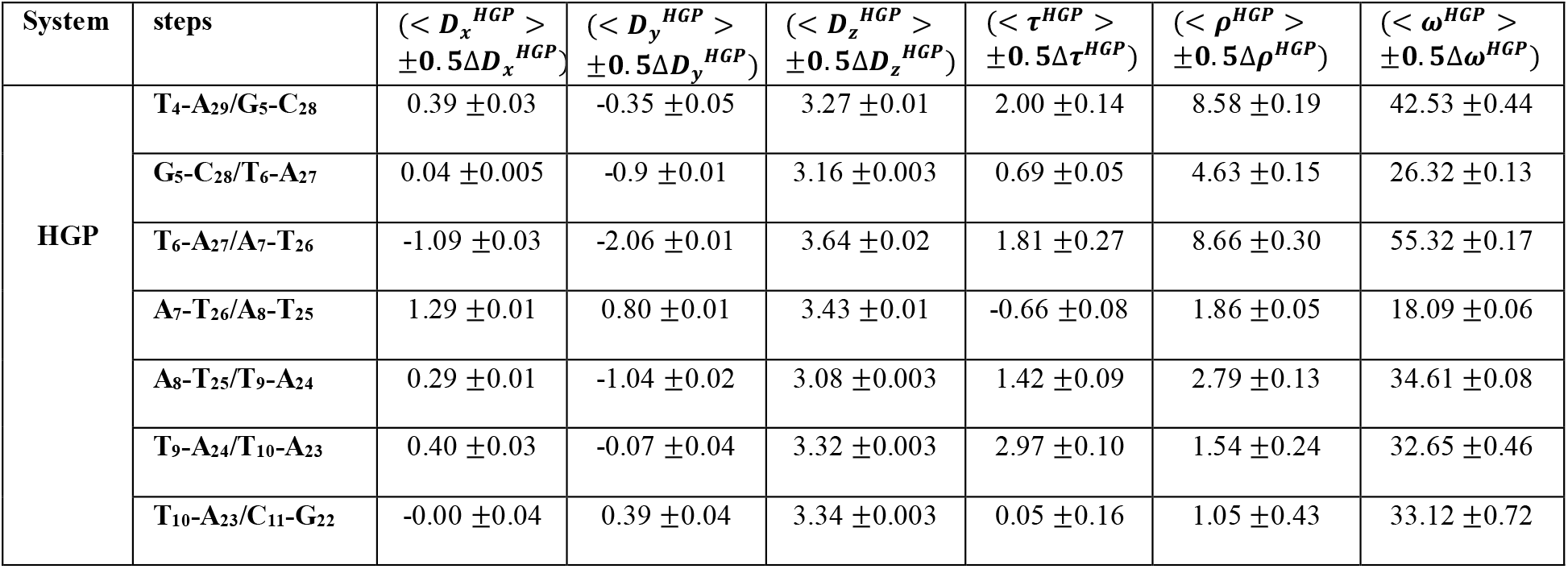
The mean with error in the mean of each step in the HGP system for the bp step parameters (D_x_, D_y_, D_z_, τ, ρ, ω). D_x_, D_y_, D_z_ are in A^0^. τ, ρ and ω are in degree.

**Table S4:**
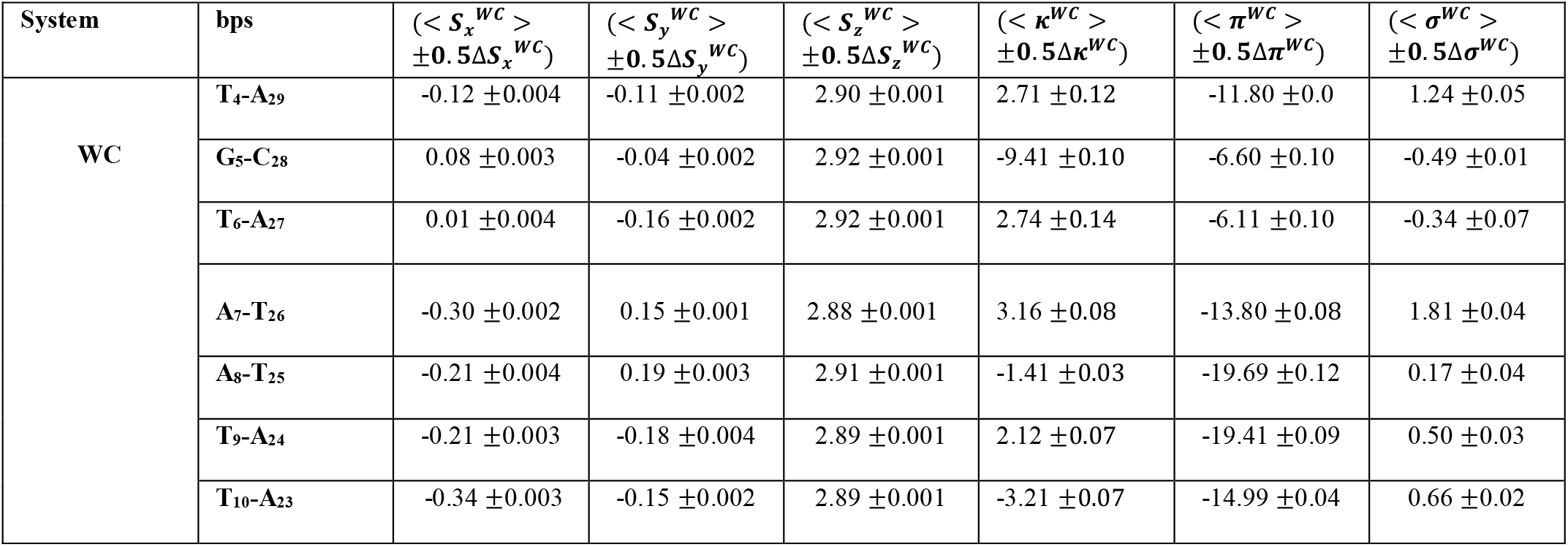
The mean and the error in the mean of each bp in the WC system for all intra bp parameters (S_x_, S_y_, S_z_, κ, π, σ). S_x_, S_y_, S_z_ are in A^0^. κ, π and σ are in degree.

**Table S5:**
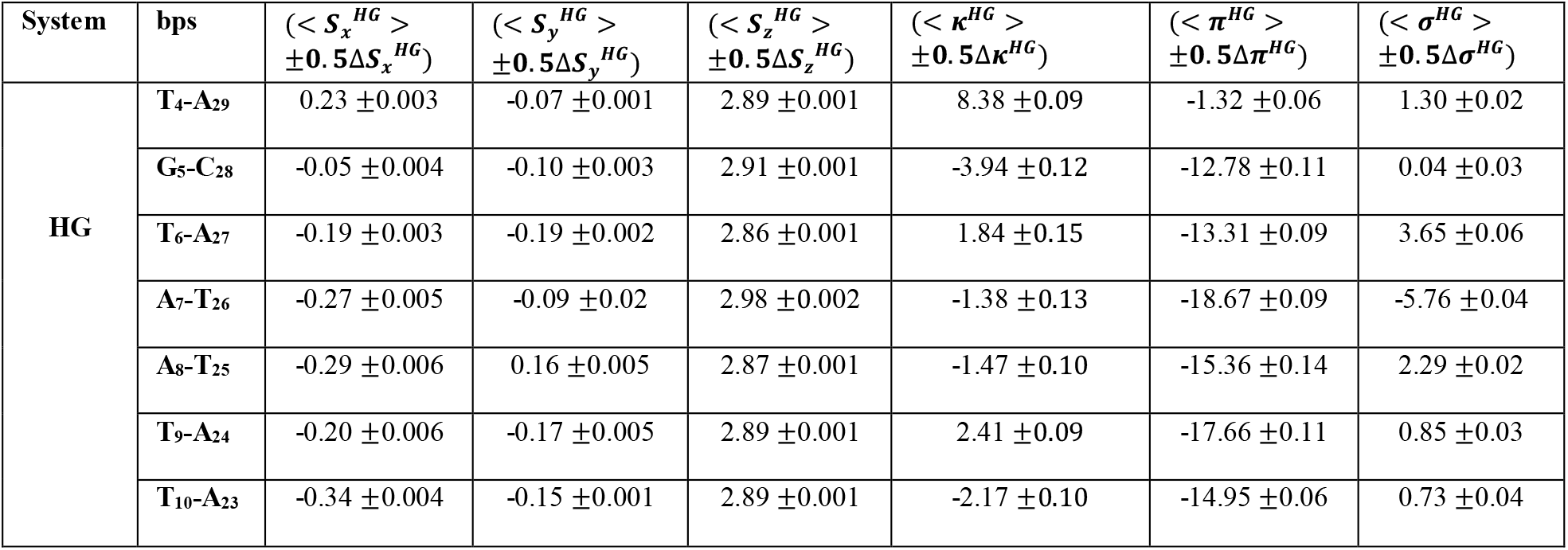
The mean and the error in the mean of each bp in the HG system for all intra bp parameters (S_x_, S_y_, S_z_, κ, π, σ). S_x_, S_y_, S_z_ are in A^0^. κ, π and σ are in degree.

| System | bps | $(\langle S_x^{HG} \rangle \pm 0.5\Delta S_x^{HG})$ | $(\langle S_y^{HG} \rangle \pm 0.5\Delta S_y^{HG})$ | $(\langle S_z^{HG} \rangle \pm 0.5\Delta S_z^{HG})$ | $(\langle \kappa^{HG} \rangle \pm 0.5\Delta \kappa^{HG})$ | $(\langle \pi^{HG} \rangle \pm 0.5\Delta \pi^{HG})$ | $(\langle \sigma^{HG} \rangle \pm 0.5\Delta \sigma^{HG})$ |
| --- | --- | --- | --- | --- | --- | --- | --- |
| HG | T4-A29 | $0.23 \pm 0.003$ | $-0.07 \pm 0.001$ | $2.89 \pm 0.001$ | $8.38 \pm 0.09$ | $-1.32 \pm 0.06$ | $1.30 \pm 0.02$ |
| | G5-C28 | $-0.05 \pm 0.004$ | $-0.10 \pm 0.003$ | $2.91 \pm 0.001$ | $-3.94 \pm 0.12$ | $-12.78 \pm 0.11$ | $0.04 \pm 0.03$ |
| | T6-A27 | $-0.19 \pm 0.003$ | $-0.19 \pm 0.002$ | $2.86 \pm 0.001$ | $1.84 \pm 0.15$ | $-13.31 \pm 0.09$ | $3.65 \pm 0.06$ |
| | A7-T26 | $-0.27 \pm 0.005$ | $-0.09 \pm 0.02$ | $2.98 \pm 0.002$ | $-1.38 \pm 0.13$ | $-18.67 \pm 0.09$ | $-5.76 \pm 0.04$ |
| | A8-T25 | $-0.29 \pm 0.006$ | $0.16 \pm 0.005$ | $2.87 \pm 0.001$ | $-1.47 \pm 0.10$ | $-15.36 \pm 0.14$ | $2.29 \pm 0.02$ |
| | T9-A24 | $-0.20 \pm 0.006$ | $-0.17 \pm 0.005$ | $2.89 \pm 0.001$ | $2.41 \pm 0.09$ | $-17.66 \pm 0.11$ | $0.85 \pm 0.03$ |
| | T10-A23 | $-0.34 \pm 0.004$ | $-0.15 \pm 0.001$ | $2.89 \pm 0.001$ | $-2.17 \pm 0.10$ | $-14.95 \pm 0.06$ | $0.73 \pm 0.04$ |

**Table S6:** The mean and the error in the mean of each bp in the HGP system for all intra bp parameters (S_x_, S_y_, S_z_, κ, π, σ). S_x_, S_y_, S_z_ are in A^0^. κ, π and σ are in degree.

| System | bps | $(\langle S_x^{HGP} \rangle \pm 0.5\Delta S_x^{HGP})$ | $(\langle S_y^{HGP} \rangle \pm 0.5\Delta S_y^{HGP})$ | $(\langle S_z^{HGP} \rangle \pm 0.5\Delta S_z^{HGP})$ | $(\langle \kappa^{HGP} \rangle \pm 0.5\Delta \kappa^{HGP})$ | $(\langle \pi^{HGP} \rangle \pm 0.5\Delta \pi^{HGP})$ | $(\langle \sigma^{HGP} \rangle \pm 0.5\Delta \sigma^{HGP})$ |
| --- | --- | --- | --- | --- | --- | --- | --- |
| HGP | T4-A29 | $0.14 \pm 0.01$ | $-0.0 \pm 0.01$ | $2.90 \pm 0.001$ | $12.00 \pm 0.20$ | $-1.87 \pm 0.22$ | $0.87 \pm 0.11$ |
| | G5-C28 | $0.01 \pm 0.01$ | $-0.48 \pm 0.02$ | $2.92 \pm 0.005$ | $-3.67 \pm 0.53$ | $-15.75 \pm 0.25$ | $2.84 \pm 0.28$ |
| | T6-A27 | $-0.27 \pm 0.02$ | $-0.11 \pm 0.005$ | $2.86 \pm 0.006$ | $-0.10 \pm 0.49$ | $-18.92 \pm 0.16$ | $11.45 \pm 0.17$ |
| | A7-T26 | $0.01 \pm 0.01$ | $-0.01 \pm 0.005$ | $2.97 \pm 0.004$ | $-16.97 \pm 0.53$ | $-11.73 \pm 0.28$ | $-6.84 \pm 0.14$ |
| | A8-T25 | $-0.18 \pm 0.01$ | $0.16 \pm 0.006$ | $2.82 \pm 0.002$ | $-13.58 \pm 0.38$ | $-20.21 \pm 0.19$ | $8.69 \pm 0.13$ |
| | T9-A24 | $-0.11 \pm 0.01$ | $-0.15 \pm 0.006$ | $2.89 \pm 0.002$ | $-9.58 \pm 0.38$ | $-18.12 \pm 0.30$ | $2.19 \pm 0.31$ |
| | T10-A23 | $-0.34 \pm 0.01$ | $-0.13 \pm 0.004$ | $2.91 \pm 0.003$ | $-5.23 \pm 0.54$ | $-11.20 \pm 0.27$ | $0.28 \pm 0.16$ |

**Table S7:** The mean and the error of the mean value of phase angle (P) of each base in the WC, HG and HGP systems.

| bases | $(\langle P^{WC} \rangle \pm 0.5\Delta P^{WC})$<br>(degree) | $(\langle P^{HG} \rangle \pm 0.5\Delta P^{HG})$<br>(degree) | $(\langle P^{HGP} \rangle \pm 0.5\Delta P^{HGP})$<br>(degree) |
| --- | --- | --- | --- |
| T<sub>4</sub> | 185.55 ± 0.06 | 182.35 ± 0.48 | 188.67 ± 1.18 |
| G<sub>5</sub> | 186.72 ± 0.09 | 182.82 ± 0.25 | 178.78 ± 0.26 |
| T<sub>6</sub> | 185.29 ± 0.12 | 181.24 ± 0.13 | 176.66 ± 0.38 |
| A<sub>7</sub> | 176.46 ± 0.06 | 222.53 ± 0.42 | 177.76 ± 0.05 |
| A<sub>8</sub> | 179.07 ± 0.07 | 179.31 ± 0.07 | 179.52 ± 0.39 |
| T<sub>9</sub> | 187.96 ± 0.21 | 188.24 ± 0.30 | 202.81 ± 0.87 |
| T<sub>10</sub> | 191.64 ± 0.27 | 189.96 ± 0.22 | 183.67 ± 0.64 |
| A<sub>23</sub> | 177.58 ± 0.03 | 177.95 ± 0.09 | 177.61 ± 0.35 |
| A<sub>24</sub> | 179.84 ± 0.06 | 181.41 ± 0.15 | 179.54 ± 0.24 |
| T<sub>25</sub> | 185.64 ± 0.31 | 185.52 ± 0.33 | 182.17 ± 0.49 |
| T<sub>26</sub> | 189.99 ± 0.51 | 201.37 ± 0.41 | 188.44 ± 0.36 |
| A<sub>27</sub> | 180.55 ± 0.13 | 178.67 ± 0.07 | 178.64 ± 0.07 |
| C<sub>28</sub> | 181.23 ± 0.16 | 181.55 ± 0.17 | 181.13 ± 0.51 |
| A<sub>29</sub> | 179.78 ± 0.05 | 182.30 ± 0.21 | 181.91 ± 0.20 |

**Table S8:** Mean and error of the mean of *d*_*D*−*A*_ and 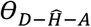 in A_7_-T_26_ bp.

| System | $\langle d_{N7-N3} \rangle$<br>$\pm 0.5\Delta d_{N7-N3}$<br>(nm) | $\langle \theta_{N7-H-N3} \rangle$<br>$\pm 0.5\Delta \theta_{N7-H-N3}$<br>(degree) | $\langle d_{N1-N3} \rangle$<br>$\pm 0.5\Delta d_{N1-N3}$<br>(nm) | $\langle \theta_{N1-H-N3} \rangle$<br>$\pm 0.5\Delta \theta_{N1-H-N3}$<br>(degree) |
| --- | --- | --- | --- | --- |
| WC | 0.64 ± 0.0001 | 159.09 ± 0.01 | 0.29 ± 0.0001 | 169.83 ± 0.02 |
| HG | 0.31 ± 0.0009 | 168.70 ± 0.14 | 0.59 ± 0.0001 | 145.19 ± 0.03 |
| HGP | 0.30 ± 0.0004 | 169.09 ± 0.04 | 0.59 ± 0.0002 | 144.67 ± 0.10 |

**Table S9:** Mean and error of the mean values, of *d*_*C*1′_ for T_6_-A_27_, A_7_-T_26_ and A_8_-T_25_ bps, and *d*_*p*7−*p*38_ and *d*_*p*8−*p*37_ for all systems.

| System | $\langle d_{C1'}^{T_6:A_{27}} \rangle$<br>$\pm 0.5\Delta d_{C1'}^{T_6:A_{27}}$<br>(nm) | $\langle d_{C1'}^{A_7:T_{26}} \rangle$<br>$\pm 0.5\Delta d_{C1'}^{A_7:T_{26}}$<br>(nm) | $\langle d_{C1'}^{A_8:T_{25}} \rangle$<br>$\pm 0.5\Delta d_{C1'}^{A_8:T_{25}}$<br>(nm) | $\langle d_{P7-P38} \rangle$<br>$\pm 0.5\Delta d_{P7-P38}$<br>(nm) | $\langle d_{P8-P37} \rangle$<br>$\pm 0.5\Delta d_{P8-P37}$<br>(nm) |
| --- | --- | --- | --- | --- | --- |
| WC | 1.05±0.001 | 1.05±0.0002 | 1.05±0.0002 | 1.76 ±0.0003 | 1.77 ±0.0002 |
| HG | 1.03±0.0004 | 0.90±0.0003 | 1.03±0.0001 | 1.70 ±0.0002 | 1.71 ±0.0004 |
| HGP | 1.00±0.001 | 0.89±0.002 | 1.00±0.001 | 1.62 ±0.003 | 1.70 ±0.002 |

**Table S10:** Mean and error of the mean value of χ for T_6_, A_7_, A_8_ and T_26_ bases.

| System | $\langle \chi_{T_6} \rangle$<br>$\pm 0.5\Delta \chi_{T_6}$<br>(degree) | $\langle \chi_{A_7} \rangle$<br>$\pm 0.5\Delta \chi_{A_7}$<br>(degree) | $\langle \chi_{A_8} \rangle$<br>$\pm 0.5\Delta \chi_{A_8}$<br>(degree) | $\langle \chi_{T_{26}} \rangle$<br>$\pm 0.5\Delta \chi_{T_{26}}$<br>(degree) |
| --- | --- | --- | --- | --- |
| WC | -111.14±0.08 | -98.47±0.13 | -112.77±0.06 | -111.24±0.19 |
| HG | -112.43±0.05 | 54.73±0.12 | -98.64±0.10 | -120.03±0.13 |
| HGP | -127.44±0.27 | 60.36±0.23 | -96.88±0.28 | -105.75±0.17 |

**Table S11:** Mean and error of the mean value of α and *γ* torsion angles in A_7_ and T_26_ nucleotides.

| System | $\langle \alpha_{A_7} \rangle \pm 0.5\Delta \alpha_{A_7}$<br>(degree) | $\langle \gamma_{A_7} \rangle \pm 0.5\Delta \gamma_{A_7}$<br>(degree) | $\langle \alpha_{T_{26}} \rangle \pm 0.5\Delta \alpha_{T_{26}}$<br>(degree) | $\langle \gamma_{T_{26}} \rangle \pm 0.5\Delta \gamma_{T_{26}}$<br>(degree) |
| --- | --- | --- | --- | --- |
| WC | -70.51 ±0.13 | 53.48 ±0.05 | -66.54 ±0.14 | 55.27 ±0.14 |
| HG | -56.51 ±0.57 | 44.14 ±0.54 | -66.80 ±0.31 | 54.41 ±0.29 |
| HGP | 63.16 ±0.23 | -65.94 ±0.22 | -77.80 ±0.21 | 59.95 ±0.12 |

**Table S12:**
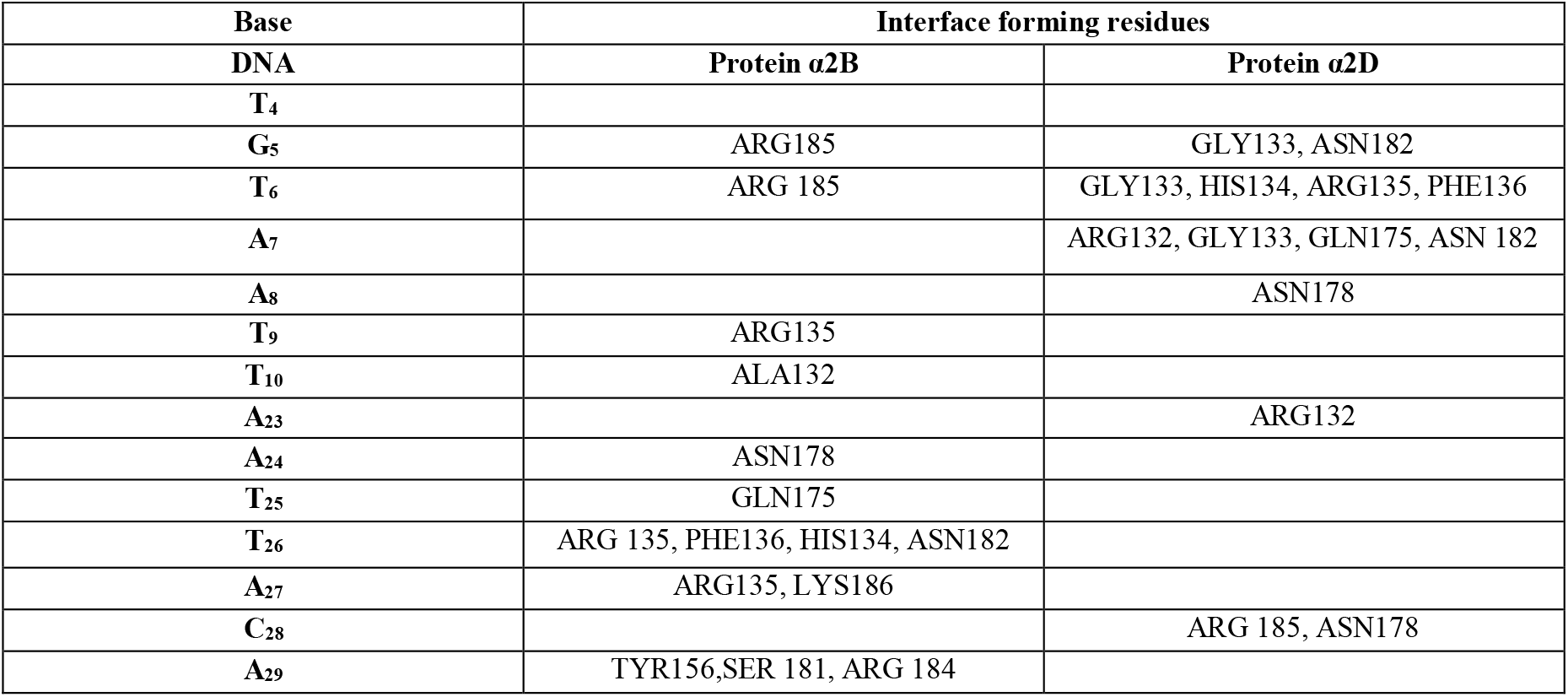
The protein residues that strongly interact with DNA bps in the HGP system.

**Table S13:** The ratio of the number of frames in each non-bonded interaction to the total number of frames, for each interface forming α2D protein residue, considering the interfacial contact between A_7_ and α2D residues.

| System | Nucleotide | Protein | H-bond | Salt bridge | Electrostatic |
| --- | --- | --- | --- | --- | --- |
| HGP | A<sub>7</sub> | $\alpha 2D$ | ARG132<br>(0.65) | ARG132<br>(0.26) | ARG132<br>(0.93) |
|  |  |  | GLY133<br>(0.92) |  |  |
|  |  |  | GLN175<br>(0.93) |  |  |

**Table S14:** The ratio of the number of frames in each non-bonded interaction to the total number of frames in the equilibrated trajectory of HGP, for each α2B protein residue, considering the interfacial contact between T_26_ and α2B residues.

| System | Nucleotide | Protein | H-bond |
| --- | --- | --- | --- |
| HGP | T<sub>26</sub> | $\alpha 2B$ | HIS134 (0.61) |
|  |  |  | PHE136 (0.99) |
|  |  |  | TRP179 (0.59) |

